# Microtubule Nucleation by Single Human γTuRC in a Partly Open Asymmetric Conformation

**DOI:** 10.1101/853218

**Authors:** Tanja Consolati, Julia Locke, Johanna Roostalu, Jayant Asthana, Wei Ming Lim, Julian Gannon, Fabrizio Martino, Alessandro Costa, Thomas Surrey

## Abstract

The γ-tubulin ring complex (γTuRC) is the major microtubule nucleator in cells. However, the mechanism of its regulation is not understood. Here, we purified human γTuRC and quantitatively characterized its nucleation properties in a TIRF microscopy-based real-time nucleation assay. We find that microtubule nucleation by γTuRC is kinetically inhibited compared to microtubule elongation. Determining the cryo-EM structure of γTuRC at 4 Å resolution reveals an asymmetric conformation with only part of the complex in a ‘closed’ conformation matching the microtubule geometry. Several factors stabilise the closed conformation. One is actin in the core of the complex and others, likely MZT1 or MZT2, line the outer perimeter of the closed part of γTuRC. The opposed side of γTuRC is in an ‘open’, nucleation-incompetent conformation, leading to a structural asymmetry, explaining the kinetic inhibition of nucleation by human γTuRC. Our data suggest possible regulatory mechanisms for microtubule nucleation by γTuRC closure.

## INTRODUCTION

Microtubule nucleation in cells is spatially and temporally controlled to ensure proper cytoskeleton function. The major nucleator is the γ-tubulin ring complex (γTuRC) in which several γ-tubulin complex proteins (GCPs) arrange 14 γ-tubulins into a helical arrangement so that they can serve as a template for microtubule nucleation (Kollman et al., 2011; Tovey and Conduit, 2018). The structure of γTuRC is best understood in budding yeast where 7 smaller ‘Y-shaped’ γTuSC complexes, each consisting of 2 γ-tubulins and one copy of GCP2 and GCP3, assemble into a conically shaped assembly upon recruitment to spindle the pole body by SPC110 (Kollman et al., 2010). A cryo-electron microscopy reconstruction of budding yeast γTuSC in complex with a SPC110 fragment at 8Å resolution revealed gaps between every second γ-tubulin in γTuRC creating a mismatch with the microtubule structure (Kollman et al., 2015). Microtubule nucleation by budding yeast γTuRC in this ‘open’ conformation could be improved 2-3fold by artificially closing these gaps through chemical crosslinking, suggesting a possible mechanism for activation of nucleation by budding yeast γTuRC (Kollman et al., 2015).

In fission yeast, filamentous fungi and metazoans, some GCP2s and GCP3s are replaced in the complex by an additional GCP4, GCP5 and GCP6 proteins and in metazoan γTuRC is a stable complex whose assembly is independent of the recruitment to target structures such as centrosomes (Farache et al., 2018; Lin et al., 2015; Murphy et al., 2001; Oegema et al., 1999; Tovey and Conduit, 2018). The exact stoichiometry and subunit order of human γTuRC is not known. Several proteins have been implicated in activating γTuRC, among which are MZT1 (GCP9) and MZT2 (GCP8), also sometimes considered core components of the metazoan complex (Hutchins et al., 2010; Kollman et al., 2011; Lin et al., 2016; Teixido-Travesa et al., 2012), the recruitment factors CDK5Rap2 (functional homologue of budding yeast Spc110) (Choi et al., 2010; Lin et al., 2014; Muroyama et al., 2016), or microtubule dynamics regulators such as the microtubule polymerase chTOG (XMAP215) or the multifunctional, catastrophe suppressing protein TPX2 (Alfaro-Aco et al., 2017; Scrofani et al., 2015; Thawani et al., 2018).

A clear understanding of the mechanisms by which the efficiency of microtubule nucleation by human γTuRC is regulated is however lacking. The kinetics of microtubule nucleation either in the absence or presence of purified γTuRC have typically been measured either by following the turbidity of suspensions of nucleating microtubules over time or by fluorescence microscopy imaging of microtubules at distinct times after mixing γTuRC with tubulin (Oegema et al., 1999; Voter and Erickson, 1984). Both types of assays have disadvantages, as they cannot distinguish easily between γTuRC-mediated and spontaneous microtubule nucleation and between microtubule nucleation and effects produced by microtubule growth and/or dynamic instability.

To overcome these limitations, we developed a microscopy-based *in vitro* nucleation assay that allows the real-time observation of the nucleation of individual microtubules by single surface-immobilized human γTuRCs and we determined the structure of the γTuRC complex by cryo-electron microscopy and single particle reconstruction. We found that human γTuRC-mediated nucleation is stochastic, highly cooperative and rather inefficient. γTuRC improved the nucleation efficiency compared to spontaneous microtubule nucleation, but templated nucleation was kinetically inhibited compared to microtubule plus end elongation. A 4 Å resolution structure of human γTuRC revealed several features that are distinctly different from the structure of budding yeast γTuRC; surprisingly, γTuRC harbours actin in its central core and the complex is markedly asymmetric only partially matching the geometry of the active form of yeast γTuRC. Our results provide an explanation for the observed inefficient nucleation by human and γTuRC and suggest a possible stimulatory function of additional factors that would morph γTuRC into a fully activated form.

## RESULTS

### Purification of biotinylated human γTuRC

We generated a HeLa Kyoto cell line that expressed a biotin acceptor peptide (BAP) and monomeric blue fluorescence protein (mBFP)-tagged GCP2 from a randomly integrated gene. Tagged GCP2 became incorporated into the human γTuRC complex and was biotinylated without compromising γTuRC function as indicated by the correct localization of the fluorescent complex to centrosomes and normal cell growth. We purified ∼ 0.1 mg tagged γTuRC from 120 g of cells in a one-day procedure using anion exchange, biotin affinity and size exclusion chromatography (Methods) (Fig. 1a, b, Fig. S1).

**Figure 1.**
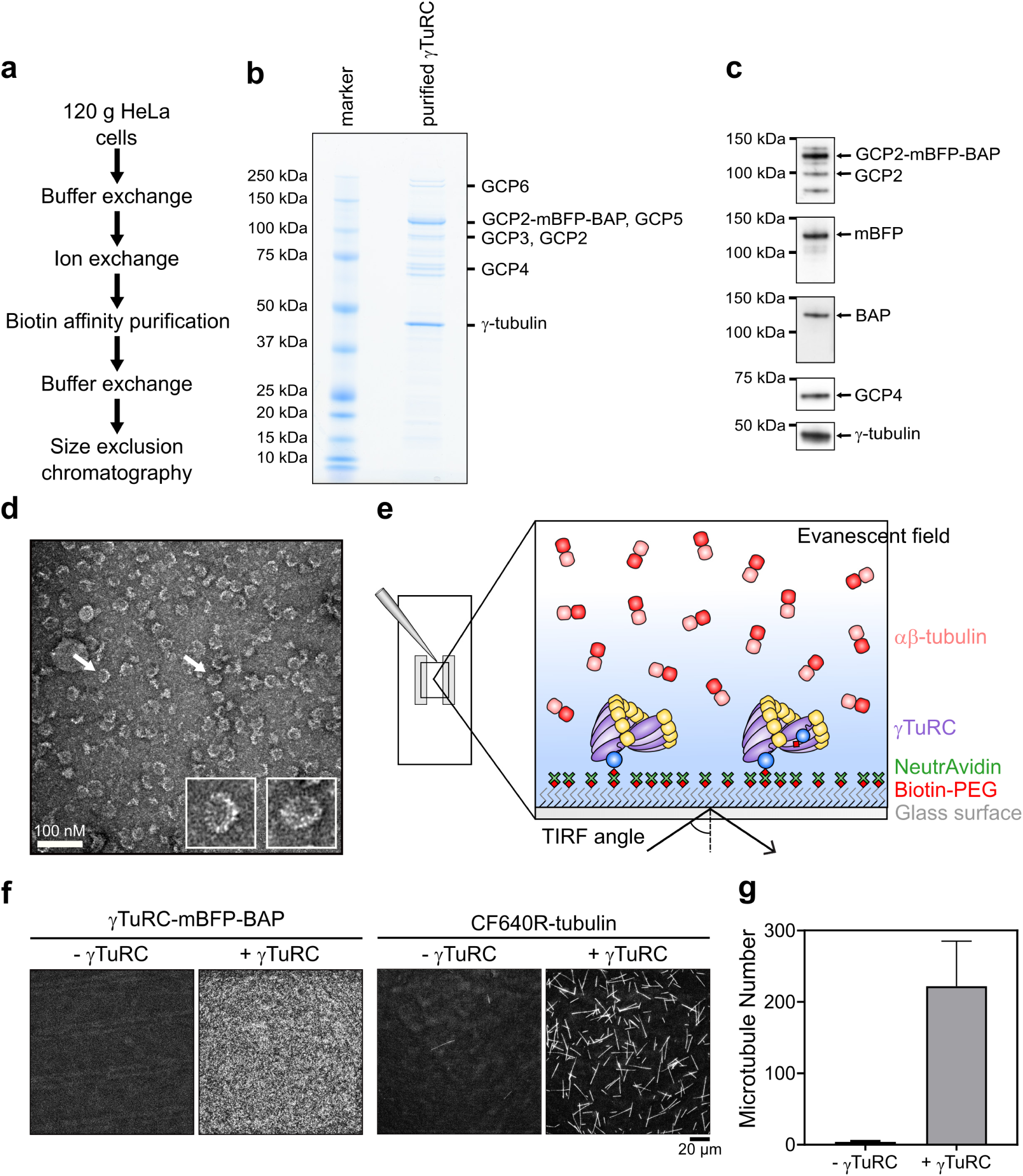
Purification and characterization of γTuRC-mBFP-BAP. **(a)** Overview of purification steps. **(b)** Coomassie-stained SDS-PAGE of purified γTuRC. Protein bands corresponding to γTuRC subunits as identified by mass spectrometry are labelled. **(c)** Western blots of purified γTuRC using antibodies against γ-tubulin, GCP2, GCP4 and mBFP. Biotinylation of the biotin acceptor peptide (BAP) was assessed by immunoblotting using horse radish peroxidase (HRP)-coupled streptavidin. **(d)** Negative-stain electron microscopy of purified γTuRC showing the expected 25-nm diameter ring structures. Two examples (white arrows) are shown as insets at higher magnification. **(e)** Schematic of total internal reflection fluorescence microscopy (TIRFM) based real-time γTuRC nucleation assay. Biotinylated fluorescent γTuRC is immobilized on a biotin-PEG-functionalized glass surface via NeutrAvidin. αβ-tubulin is added to initiate microtubule nucleation by immobilized γTuRC. **(f)** Representative TIRFM images showing the mBFP channel to visualize γTuRC on the surface (left panel) and showing the CF640R-tubulin channel to visualize nucleated microtubules (right panel) at t=20 min after start of microtubule nucleation by a temperature jump to 33 °C. 373 pM γTuRC was used for immobilization and the final CF640R-tubulin concentration was 15 μM. A representative control without γTuRC is also shown. Intensities in the images are directly comparable. **(g)** Bar graph of the average microtubule number nucleated by surface immobilized γTuRC (373 pM) within 15 min in presence of 15 μM CF640R-tubulin (n=3). Error bars are s.d. Scale bars as indicated. t=0 is 2 min after placing the sample at 33 °C. See also Fig. S1, Fig. S2 and Fig. S3.

Using mass spectroscopy, we identified all human γTuRC core subunits (γ-tubulin, GCP2-6), as well as MZT1, MZT2 and actin, which were co-purified in previous γTuRC purifications (Fig. S2 and Supplementary Material) (Choi et al., 2010; Hutchins et al., 2010; Oegema et al., 1999; Teixido-Travesa et al., 2012; Thawani et al., 2018). Additionally, we verified most subunits also by western blot using specific antibodies (Fig. 1c), indicating that the complete human γTuRC complex was successfully purified using our biotin affinity purification strategy. Purified γTuRC appeared as the typical characteristic ‘rings’ with ∼25 nm diameter in negative-stain electron microscopy images (Fig. 1d) (Zheng et al., 1995), confirming the presence of properly assembled complexes.

### γTuRC nucleates single microtubules and caps their minus ends

Next, we set up a microscopy-based real-time γTuRC-mediated microtubule nucleation assay (Fig. 1e). We used NeutrAvidin to immobilize purified biotinylated γTuRC on a biotin-polyethylene glycol (PEG) functionalized glass surface. Specific immobilization of biotinylated and mBFP-tagged γTuRC could be verified by measuring the mBFP fluorescence on NeutrAvidin-surfaces and on surfaces lacking NeutrAvidin using total internal reflection fluorescence (TIRF) microscopy (Fig. 1f, Fig. S2). In the presence of CF640R-labelled tubulin, microtubules nucleated from the γTuRC-coated surface, whereas hardly any microtubules nucleated in the absence of γTuRC under these conditions (Fig. 1g).

Only one end of γTuRC-nucleated microtubules grew, whereas the other was tethered to the surface likely via γTuRC (Fig. 2a top rows, b left, Movie 1). In contrast, when microtubules elongated from surface-immobilized, stabilized microtubule ‘seeds’ in control experiments, both microtubule ends grew (Fig. 2a, bottom row, b, right) with the faster plus end growth speed of 26.8 nm/s essentially equating the growth speed of γTuRC-nucleated microtubules of 26.3 nm/s (Fig. 2c, top). This observation demonstrates that γTuRC-nucleated microtubules grow exclusively at their plus end. Minus ends grew from ‘seeds’ with a speed of 7 nm/s, while surface anchored minus ends of γTuRC-nucleated microtubules did not grow (Fig. 2c, bottom). mGFP-labelled growing microtubule end marker EB3 decorated only the growing plus ends of γTuRC-nucleated microtubules (Fig. 2d, e, Movie 2), similar to the situation in the cell (Akhmanova and Steinmetz, 2015). In rare cases, γTuRC-nucleated microtubules first grew out of the TIRF field and later, remaining γTuRC-anchored, aligned with the surface only growing with one end (Fig. S4a, b). These data clearly demonstrate that, at our experimental conditions, plus ends of γTuRC-nucleated microtubules are dynamic and minus ends are capped by γTuRC. The occasional microtubule that nucleated in solution and then landed on the glass surface was easily distinguished from γTuRC-nucleated microtubules, as it became suddenly visible as an already elongated microtubule, displaying two dynamic microtubule ends and often also diffusing along the surface (Fig. S4c, d).

**Figure 2.**
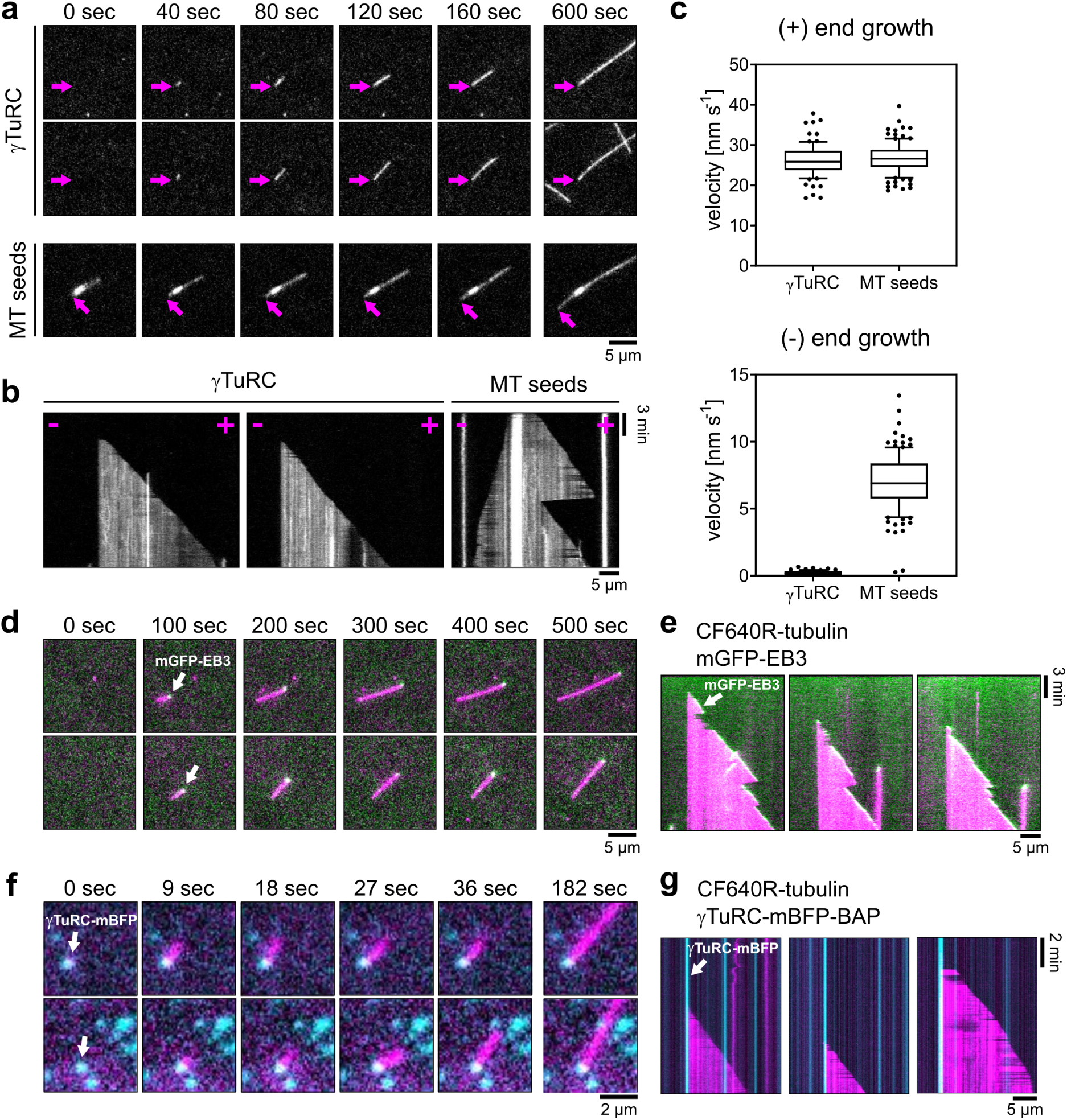
γTuRC nucleates and caps microtubules at their minus-end. **(a)** Representative time series of individual microtubules (2 top rows of panels) nucleated on a γTuRC surface (373 pM γTuRC used for immobilization) in the presence of 15 μM CF640R-tubulin showing a static (purple arrow) and a fast growing microtubule end. A control without γTuRC (bottom row) shows a microtubule growing from a stabilized microtubule ‘seed’ at the same tubulin concentration, displaying two growing microtubule ends with the minus-end (purple arrow) growing more slowly than the plus-end. **(b)** Box-and-whiskers plots of microtubule plus-end (top) and minus-end (bottom) growth speeds for γTuRC nucleation assays and microtubule seed assays (assay conditions as in (a)). Data were pooled from two data sets. Number of observed microtubule growth episodes per conditions: γTuRC nucleation assay, plus-end growth: n=86, minus-end growth: n=71; microtubule seed assay, plus end growth: n=110, minus-end growth: n=123. For the box-and-whiskers plots, boxes range from 25^th^ to 75^th^ percentile, the whiskers span from 10^th^ to 90^th^ percentile, and the horizontal line marks the mean value. **(c)** Representative TIRFM kymographs of microtubules nucleated by surface-immobilized γTuRC. For comparison a kymograph of a microtubule grown from a microtubule seed at the same tubulin concentration (as in (a)) is shown. **(d)** Representative time series of merged TIRFM images of two individual microtubules nucleated from a γTuRC surface (373 pM) in the presence of 12.5 μM CF640R-tubulin (magenta) and 200 nM mGFP-EB3 (green). mGFP-EB3 tracks the growing microtubule plus-end (white arrow), while the microtubule minus-end is static. **(e)** Corresponding TIRFM kymographs. **(f)** Representative time series of merged TIRFM images of individual microtubules nucleated from single immobilized γTuRC molecules (cyan, white arrow, 27 pM γTuRC used for immobilization) in the presence of 20 μM CF640R-tubulin (magenta). **(g)** Corresponding TIRFM kymographs. Scale bars as indicated. t=0 is 2 min after placing the sample at 33 °C. See also Fig. S4, Movie 1, Movie 2 and Movie 3.

Imaging the mBFP fluorescence of single immobilized γTuRC complexes at a reduced γTuRC density revealed that all surface-nucleated microtubules originated from a mBFP-labelled γTuRC (Fig. 2f, g, Movie 3). Taken together, these data demonstrate that immobilized γTuRC stimulates microtubule nucleation, generating microtubules with a capped minus and a dynamic plus end. We did not observe microtubule detachment from immobilized γTuRC, indicating that γTuRC is stably bound to its nucleated microtubule within the entire duration of our experiments (20 min).

### γTuRC-mediated microtubule nucleation is stochastic, cooperative and kinetically inhibited

Next, we quantified the number of γTuRC-nucleated microtubules per field of view, excluding the small fraction of microtubules nucleated in solution. Counting the γTuRC-nucleated microtubules, showed a linear increase of their number with time (Fig. 3a, b top left, Movie 4). This demonstrates that γTuRC-mediated nucleation is a stochastic process with constant nucleation probability. Increasing the γTuRC density on the surface while keeping the tubulin concentration constant at 15 μM, demonstrated that the overall nucleation rate (increase of microtubule number per time and per surface area) increased with the γTuRC concentration used for γTuRC surface immobilization (Fig. 3a, b top right, Movie 4) and with the measured mBFP intensity at the surface, i.e. the γTuRC density (Fig. 3b bottom right). The microtubule growth speed was unaffected by the γTuRC density (Fig 3b bottom left), in agreement with the tubulin concentration remaining unchanged in these experiments. We conclude that γTuRC stimulates nucleation in a dose-dependent manner.

**Figure 3.**
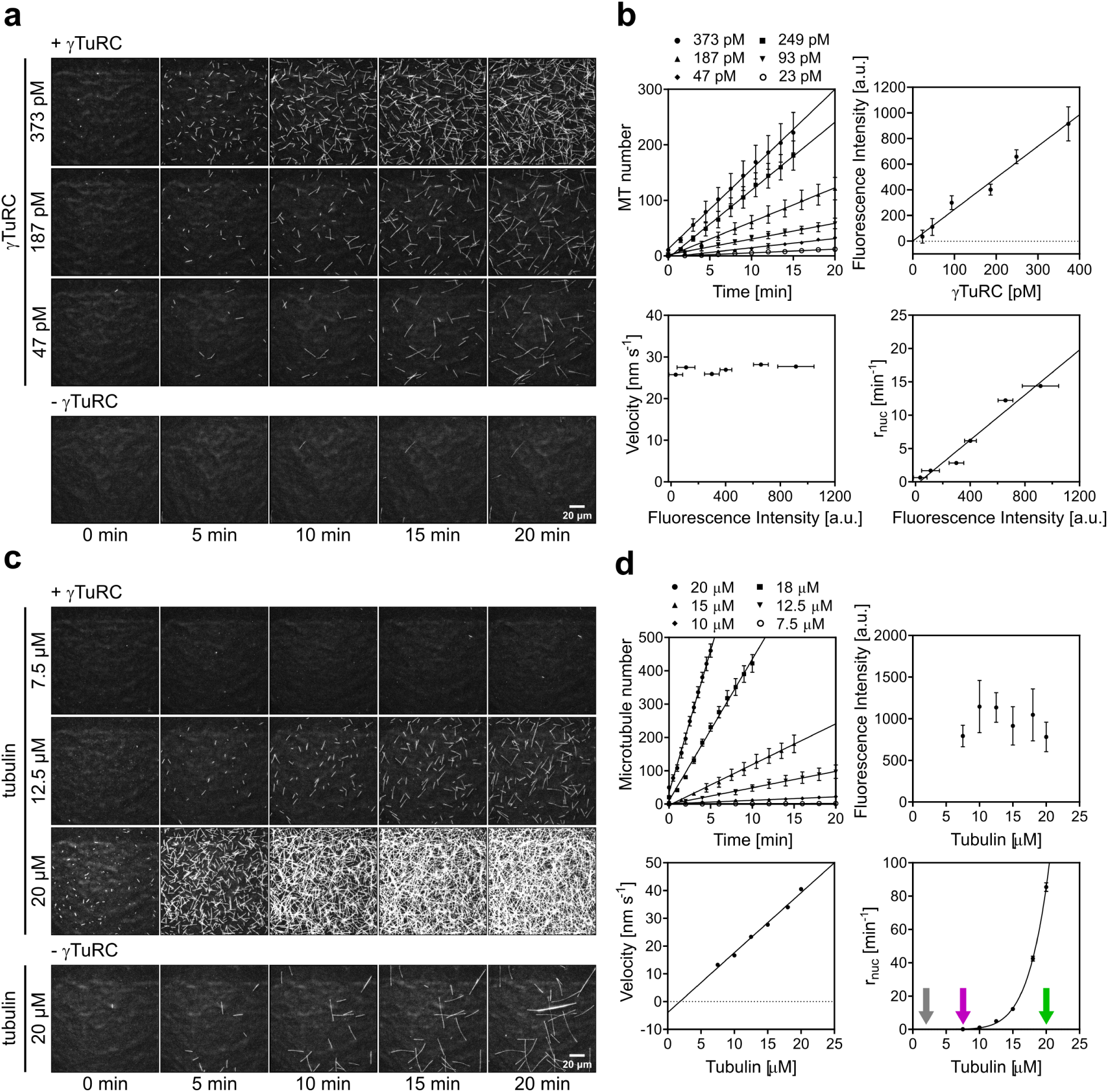
The microtubule nucleation efficiency of γTuRC depends on γTuRC surface density and tubulin concentration. **(a)** Representative time series of TIRFM images of microtubule nucleation in the presence of 15 μM CF640R-tubulin at the indicated γTuRC concentrations used for immobilization (top panel). For comparison, spontaneous microtubule nucleation in the absence of γTuRC at the same tubulin concentration is shown (bottom panel). **(b)** Plots showing a linear increase in microtubule number over time (top left), the mean γTuRC surface density (mBFP fluorescence in the field of view, top right), the mean microtubule plus end growth speed (bottom left) and the mean microtubule nucleation rate (bottom right) at different γTuRC concentrations used for immobilization. Data were pooled from at least three data sets. Number of observed microtubule growth episodes per conditions: 23 pM, n=27; 47 pM, n=64; 93 pM, n=96; 187 pM, n=191; 249 pM, n=160; 373 pM, n=302. Lines represent the linear regression. Nucleation rates (r_nuc_) were taken from the slope of the linear regression of the increase of microtubule number over time. **(c)** Representative time series of TIRFM images of microtubule nucleation in the presence of the different CF640R-tubulin concentrations, as indicated, from a γTuRC surface (373 pM) (top panel). For comparison, spontaneous microtubule nucleation in the absence of γTuRC is shown for the highest tested tubulin concentration (20 μM) (bottom panel). Spontaneous microtubule nucleation was always much less than γTuRC-mediated nucleation comparing the same tubulin concentrations (not shown). **(d)** Plots showing the linear increase in microtubule number over time (top left), the mean γTuRC surface density (mBFP fluorescence in the field of view, top right), the mean microtubule plus end growth speed (bottom left) and the mean microtubule nucleation rate (bottom right) at different tubulin concentrations. Critical tubulin concentration for microtubule elongation (grey), γTuRC-mediated nucleation (purple) and spontaneous nucleation in the absence of γTuRC (green) are marked with arrows. Data were pooled from at least three data sets. Number of observed microtubule growth episodes per conditions: 7.5 μM, n=9; 10 μM, n=51; 12.5 μM, n=210; 15 μM, n=190; 18 μM, n=237; 20 μM, n=244. The plot of the nucleation rate against tubulin concentration was fit using a power law function. All other lines represent a linear regression. All error bars are s.e.m. Field of view was always 164 μm x 164 μm. a. u., arbitrary units. Fluorescence intensities are directly comparable. Scale bars as indicated. t=0 is 2 min after placing the sample at 33 °C. See also Movie 4 and Movie 5.

Next, we changed the tubulin concentration keeping the γTuRC density constant (Fig. 3c, d top right, Movie 5). While the microtubule growth speed increased linearly with tubulin concentration, as expected (Fig. 3d, bottom left), the nucleation rate increased very non-linearly with tubulin concentration (Fig. 3c, d top left & bottom right). This observation matches earlier reports on spontaneous microtubule nucleation (Erickson and Pantaloni, 1981; Voter and Erickson, 1984) and on microtubule nucleation in the presence of *Xenopus γ*TuRC (Zheng et al., 1995). A fit to the γTuRC-mediated dependence of the nucleation rate on the tubulin concentration using a power law (line in Fig. 3d, bottom right) yielded an exponent of 6.7, demonstrating that γTuRC-mediated nucleation is a highly cooperative process, similar to spontaneous nucleation in solution (Fygenson et al., 1995; Voter and Erickson, 1984). The exponent also provides an estimate for the minimal size of a templated tubulin assembly on γTuRC that allows stable microtubule outgrowth. The apparent critical tubulin concentration required for γTuRC-mediated nucleation was 7.5 *μ*M and hence significantly lower than the ∼ 20 *μ*M for spontaneous nucleation in solution in the absence of γTuRC (Fig. 3d, bottom right: microtubule elongation (grey arrow), γTuRC-mediated nucleation (purple arrow), spontaneous nucleation (green arrow) (Brouhard and Rice, 2018; Gard and Kirschner, 1987a; Roostalu and Surrey, 2017). However, γTuRC-mediated nucleation required a tubulin concentration still higher than the 2 *μ*M tubulin threshold above which pre-existing microtubule plus ends elongate (Fig. 3d bottom left) (Voter and Erickson, 1984; Wieczorek et al., 2015). This indicates that templating a new microtubule from γTuRC is easier than forming a new microtubule in solution, but it is clearly kinetically inhibited compared to elongating an existing growing microtubule end.

Comparing the number of γTuRC complexes on the surface as measured by the number of mBFP dots with the number of nucleation events at 20 μM tubulin revealed that only ∼ 0.5% of complexes nucleated a microtubule within 9 min at our conditions. Therefore, γTuRC-mediated microtubule nucleation appears to be rather inefficient, suggesting that most likely additional factors are required for activating the complex, or for promoting nucleation by stabilizing a freshly nucleated nascent microtubule. Therefore, using our nucleation assay we tested the effects on human γTuRC-mediated microtubule nucleation elicited by three proteins that are known to affect microtubule dynamics by preferentially binding to microtubule ends and that have also been reported to affect nucleation in different ways.

### chTOG and TPX2 stimulate γTuRC-mediated microtubule nucleation

The microtubule polymerase chTOG/XMAP215 (Brouhard et al., 2008; Gard and Kirschner, 1987b) is known to mildly stimulate spontaneous microtubule nucleation (Ghosh et al., 2013; Roostalu et al., 2015). In the presence of human γTuRC, nucleation from surface-immobilized γTuRC is strongly promoted (Fig. 4a-c, Movie 6), in agreement with previous reports using the budding yeast and *Xenopus* orthologues of these proteins (Gunzelmann et al., 2018; Thawani et al., 2018). We find here that chTOG stimulates microtubule nucleation by human γTuRC by a factor of up to 21-fold, with the stimulatory effect saturating in a physiological chTOG concentration range (Fig. 4c). Saturation of the acceleration of microtubule plus end growth by chTOG occurs at a similar concentration (Figs. 4d, Fig. S5), suggesting that both chTOG effects are related and saturate when plus end binding sites at microtubule ends are fully occupied by chTOG. Thus, acceleration of outgrowth of a nascent microtubule forming on γTuRC may be one mechanism to increase overall nucleation efficiency (Roostalu and Surrey, 2017).

**Figure 4.**
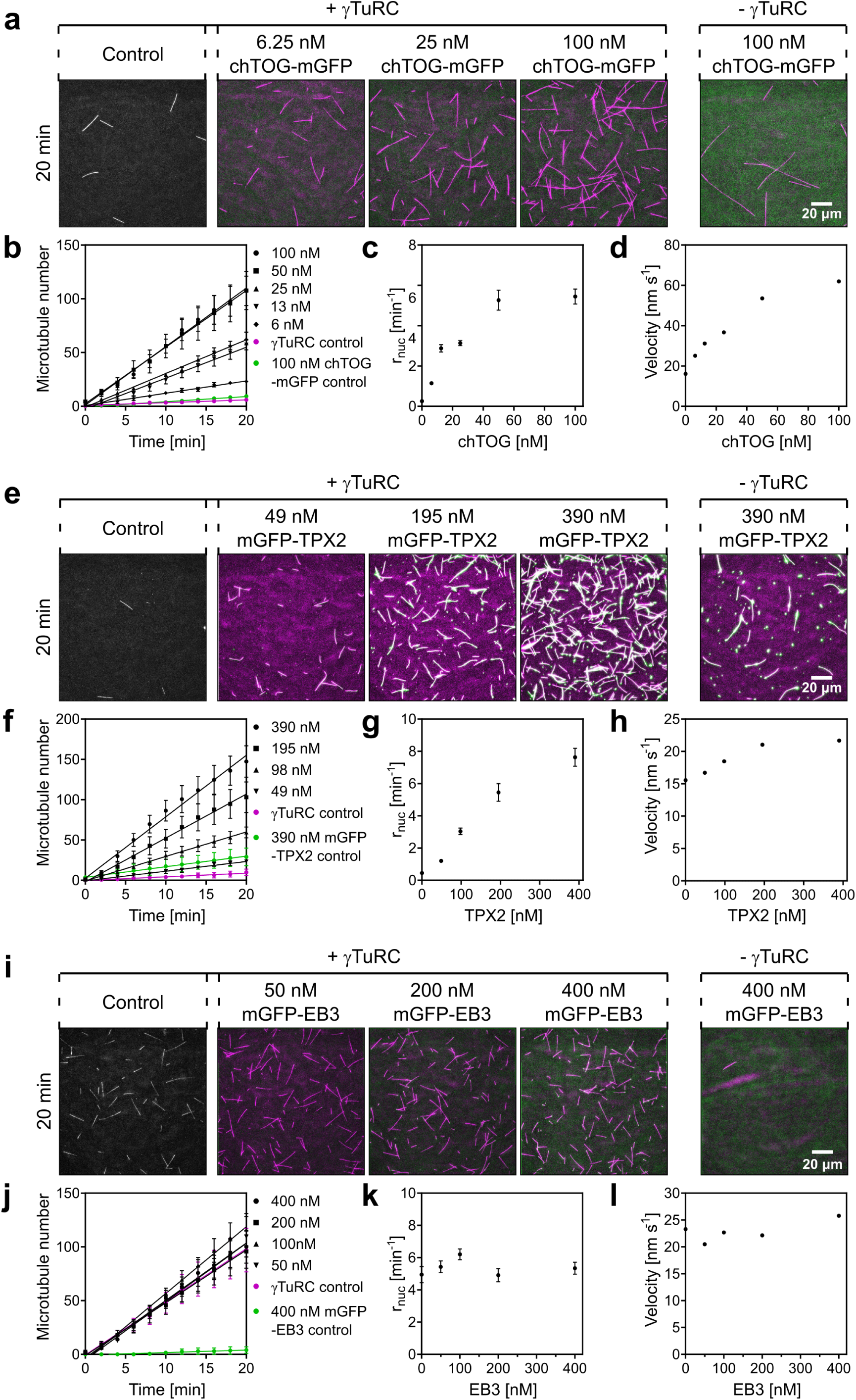
Microtubule associated proteins can increase the microtubule nucleation efficiency of γTuRC. **(a)** Representative TIRFM images of γTuRC-mediated microtubule nucleation in the presence of different chTOG-mGFP concentrations, as indicated. Microtubules are magenta, chTOG-mGFP is green. Plots showing, **(b)** linearly increasing microtubule numbers over time, **(c)** the mean microtubule nucleation rate and, **(d)** the mean microtubule growth speed at different chTOG-mGFP concentrations. Data were pooled from at least three data sets. Number of observed microtubule growth episodes per conditions: 0 nM, n=16; 6 nM, n=49; 13 nM, n=138; 25 nM, n=122; 50 nM, n=160; 100 nM, n=193. **(e)** Representative TIRFM images of γTuRC-mediated microtubule nucleation in the presence of mGFP-TPX2 concentrations (green), as indicated. Plots showing, **(f)** linearly increasing microtubule numbers over time**, (g)** the mean microtubule nucleation rate and**, (h)** the mean microtubule growth speed at different mGFP-TPX2 concentrations. Data were pooled from at least three data sets. Number of observed microtubule growth episodes per conditions: 0 nM, n=15; 49 nM, n=50; 98 nM, n=77; 195 nM, n=143; 390 nM, n=105. **(i)** Representative merged TIRFM images of γTuRC-mediated microtubule nucleation in the presence of different mGFP-EB3 concentrations (green), as indicated. Plots showing, **(j)** linearly increasing microtubule numbers over time, **(k)** the mean microtubule nucleation rate and, **(l)** the mean microtubule growth speed at different mGFP-EB3 concentrations. Data were pooled from at least three data sets. Number of observed microtubule growth episodes per conditions: 0 nM, n=210; 50 nM, n=132; 100 nM, n=230; 200 nM, n=191; 400 nM, n=290. 373 pM γTuRC was used for immobilization. chTOG and TPX2 assays were performed at 10 μM CF640R-tubulin, EB3 experiments at 12.5 μM CF640R-tubulin. As controls, representative TIRFM images in either the absence of microtubule associated proteins or absence of γTuRC are shown for the highest tested concentration of the various microtubule associated proteins. Straight lines represent linear regressions. All error bars are s.e.m. Fluorescence intensities are directly comparable. Scale bars as indicated. All TIRFM images were taken at t=20 min. t=0 is 2 min after placing the sample at 33 °C. See also Fig. S5, Fig. S6, Movie 6 and Movie 7.

TPX2, a protein involved in chromatin-dependent microtubule nucleation, can also stimulate nucleation *in vitro* (Alfaro-Aco et al., 2017; Roostalu et al., 2015; Schatz et al., 2003). Its ability to form phase-separated condensates has recently been reported to promote this activity (King and Petry, 2019). We tested here to which extent TPX2 could stimulate microtubule nucleation from immobilized γTuRC. We measured microtubules nucleating from the γTuRC surface, excluding the minor fraction of those microtubules that nucleate from TPX2 condensates (Fig. S6). We observed that TPX2 stimulated γTuRC-mediated microtubule nucleation in our assay in a dose-dependent manner, however only at rather high concentrations compared to physiological TPX2 concentrations (Fig. 4e-g, Movie 7). TPX2 hardly affected microtubule growth speed (Fig. 4h), as reported previously (Roostalu et al., 2015; Wieczorek et al., 2015). Therefore, TPX2 likely promotes γTuRC-mediated nucleation *in vitro* by a different mechanism compared to chTOG, possibly by suppressing depolymerisation of a nascent microtubule on γTuRC, through its catastrophe suppressing activity (Roostalu et al., 2015; Wieczorek et al., 2015). In contrast to chTOG and TPX2, we did not observe any effect of the plus end tracking protein EB3 on γTuRC-mediated nucleation (Fig. 4i-l).

### The cryo-EM structure of human γTuRC reveals a partly open conformation

To understand the molecular basis of microtubule nucleation, we determined the cryo-EM structure of γTuRC to a resolution of 4 Å (Fig. S7). The complex is arranged in a left-handed spiral that narrows at one end, reminiscent of a fish-and-chips newspaper cone, with a 300 Å largest diameter and a height of 200 Å (Fig. 5a). The spiral is formed by 14 similar modules (“stalks”), which support 14 globular features decorating the largest face of the complex. Comparison with the crystal structure of the GCP4 subunit of γTuRC (PDB entry 3RIP) (Guillet et al., 2011) allows the immediate identification of 14 different GCP protomers forming the spiral, which we number starting from the narrow bottom to the large-face top of the spiral. Notably, subunit 14 at the top of the spiral aligns with the lowermost subunit 1 (Fig. 5b). Inter-GCP interactions closely resemble those observed at the GCP2-GCP3 interface visible in the yeast γTuSC complex structure (Kollman et al., 2015). While GCP subunits 1 through 8 in γTuRC engage in tight inter-protomer interactions that involve both the constricted side and the larger side of the cone, subunits 9 through 14 merely interact at the tip of the cone, and appear more flexible as they depart radially from the core of the structure (Fig. 5a). The globular densities decorating the wider side of the GCP cone was rigid-body fitted by 14 γ-tubulin protomers (PDB entry 1Z5V) (Aldaz et al., 2005), resulting in a configuration akin to the γTuSC structure (Kollman et al., 2015) (Fig. 5b).

**Figure 5.**
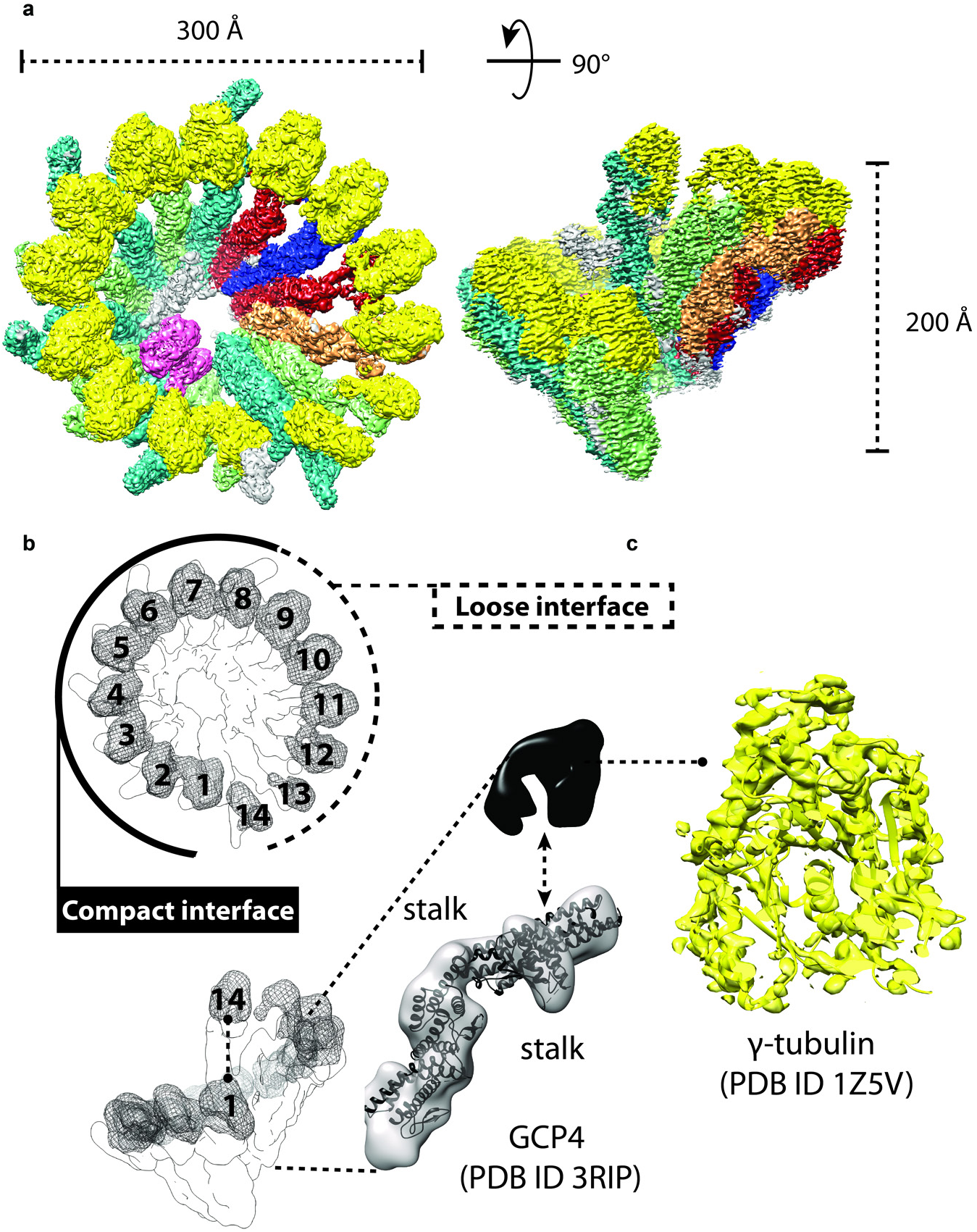
Cryo-EM structure of human γTuRC. **(a)** Surface rendering of the cryo-EM structure viewed from the top and side. γTuRC is shaped like a cone with a base diameter of 300 Å and height of 200 Å. **(b)** γTuRC contains 14 stalk protomers that support 14 globular densities. Subunits in the lowermost position 1 and the uppermost position 14 are aligned. Docking of the human GCP4 crystal structure (PDB entry 3RIP) into any of the cone positions reveals that the 14 stalk densities correspond to GCP proteins. The 14 globular densities instead correspond to γ-tubulin, as revealed by atomic docking of PDB entry 1Z5V. See also Fig. S7.

As our mass spectrometry analysis of the purified human γTuRC identified all five paralogous GCP2, 3, 4, 5 and 6 subunits, we sought to identify each subunit in our 14-mer complex. Homology searches of human GCP2 and GCP3 yielded atomic models that were very similar, although GCP3 presented a characteristic helical extension in the C-terminal, γ-tubulin interacting domain (also known as GRIP2 domain (Guillet et al., 2011; Gunawardane et al., 2000; Murphy et al., 2001)) (Fig. 6a). Given their structural homology, unique, structured sequence insertion and the well documented ability to heterodimerise, we were able to assign the GCP2 subunit to spiral protomers in position 1, 3, 5, 7, 13 and GCP3 subunit in position 2, 4, 6, 8 and 14 (Fig. 6a). To locate GCP4 in the cryo-EM map, we employed cross-correlation searches in the constricted region of the GCP spiral, where local resolution ranges from 3 to 3.5 Å. The human GCP4 N-terminal domain (“GRIP1” domain (Guillet et al., 2011)) extracted from the crystal structure showed the highest correlation at GCP in position 9 and 11. Amino acidic side chains of alpha helices in the crystallographic model match the density features in the N-terminal GCP4 cryo-EM map without the need of any real space refinement, providing us with confidence in the subunit assignment. The rest of the atomic structure was split in two additional domains (“MID” and “C-terminal”), which were docked as independent rigid bodies to achieve the best fitting results (Fig. 6b). Having assigned GCP2, 3 and 4 only two protomers in the GCP spiral remained unassigned, corresponding to positions 10 and 12. GCP5 and GCP6 both contain a characteristic internal insertion within the MID domain, although the sequence inserted in GCP6 is significantly larger. As the unoccupied cryo-EM density around the mid domain in GCP position 12 is more extensive, we tentatively assign this subunit to GCP6 (Fig. 6c, d). GCP6 also contains an extensive N-terminal extension that matches the observed cryo-EM density (Fig. 6d).

**Figure 6.**
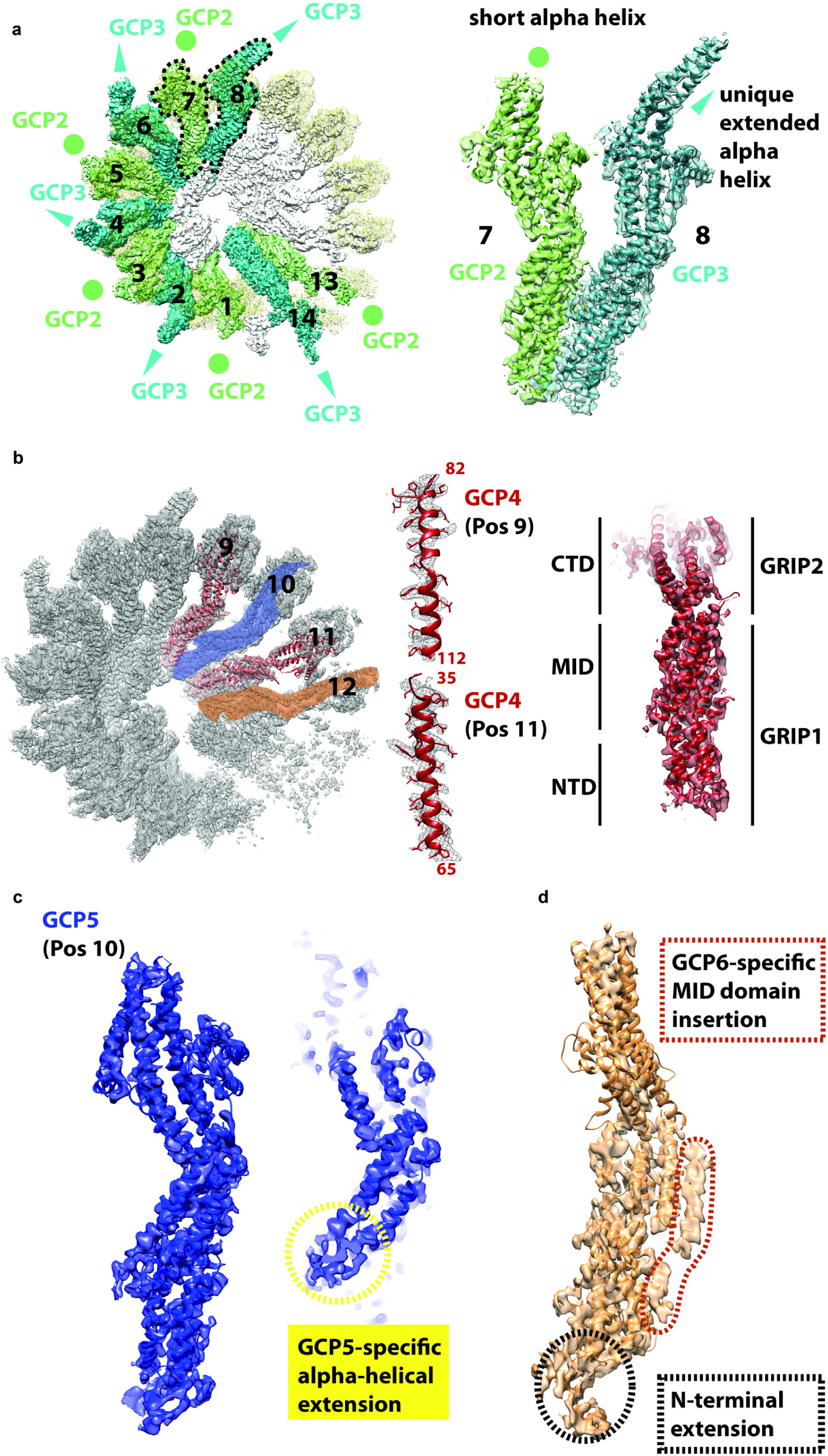
GCP subunit assignment. **(a)** GCP2 and GCP3 are known to form a stable heterodimer. Homology modelling indicates that GCP3 contains a unique *α*-helical extension, resulting in a distinctive feature that radially departs from the GCP spiral structure. This structural feature allows us to assign GCP2 and GCP3 around the γTuRC complex. **(b)** GCP4 can be assign by docking the human crystal structure into the cryo-EM map. This fitting exercise, even in the absence of any further real-space refinement, allows us to appreciate the match between amino acidic side chains from the X-ray model and the density features in the cryo-EM map. We therefore assign GCP4 to positions 9 and 11 in the map. Rigid body docking of N-terminal (GRIP1) and MID/C-terminal domains (GRIP2) is required to optimise the fitting of each individual structure. **(c)** GCP5 is assigned to position 10 due to the recognition of a characteristic predicted helical extension in the MID domain. **(d)** Assigned to position 12, GCP6 contains the largest N-terminal extension (marked in black) and MID domain insertion (marked in red) amongst GCP protomers.

### Peripherally bound MZT and an internal actin stabilize the closed conformation

Having located 14 GCP and 14 γ-tubulin protomers in the cryo-EM map, we then focused on the unassigned density. We noted that specific N-terminal GCP interfaces (in positions 1-2, 3-4, 5-6, 7-8 and 13-14) contacted a discernible *Λ*-shaped *α*-helical feature (hereafter, AHF), lining the outer perimeter of the constricted cone end. This feature appears to seal off the GCP2-GCP3 interface, occupying a position that matches that of Spc110, required for stable γTuSC complex formation in yeast (Kollman et al., 2015) (Fig. 7a). We speculated that γTuRC accessory factors such as MZT1 or MZT2, which were co-purified in our preparation, might occupy this density feature and play a complex-stabilising role similar to the yeast specific Spc110. On the C-terminal end of the spiral, we could additionally observe some unassigned C-terminal density contacting the GCP3 C-terminus, which might function as a cap that blocks further GCP polymerization, helping to define subunit composition in the γTuRC complex (Fig. 7b).

**Figure 7.**
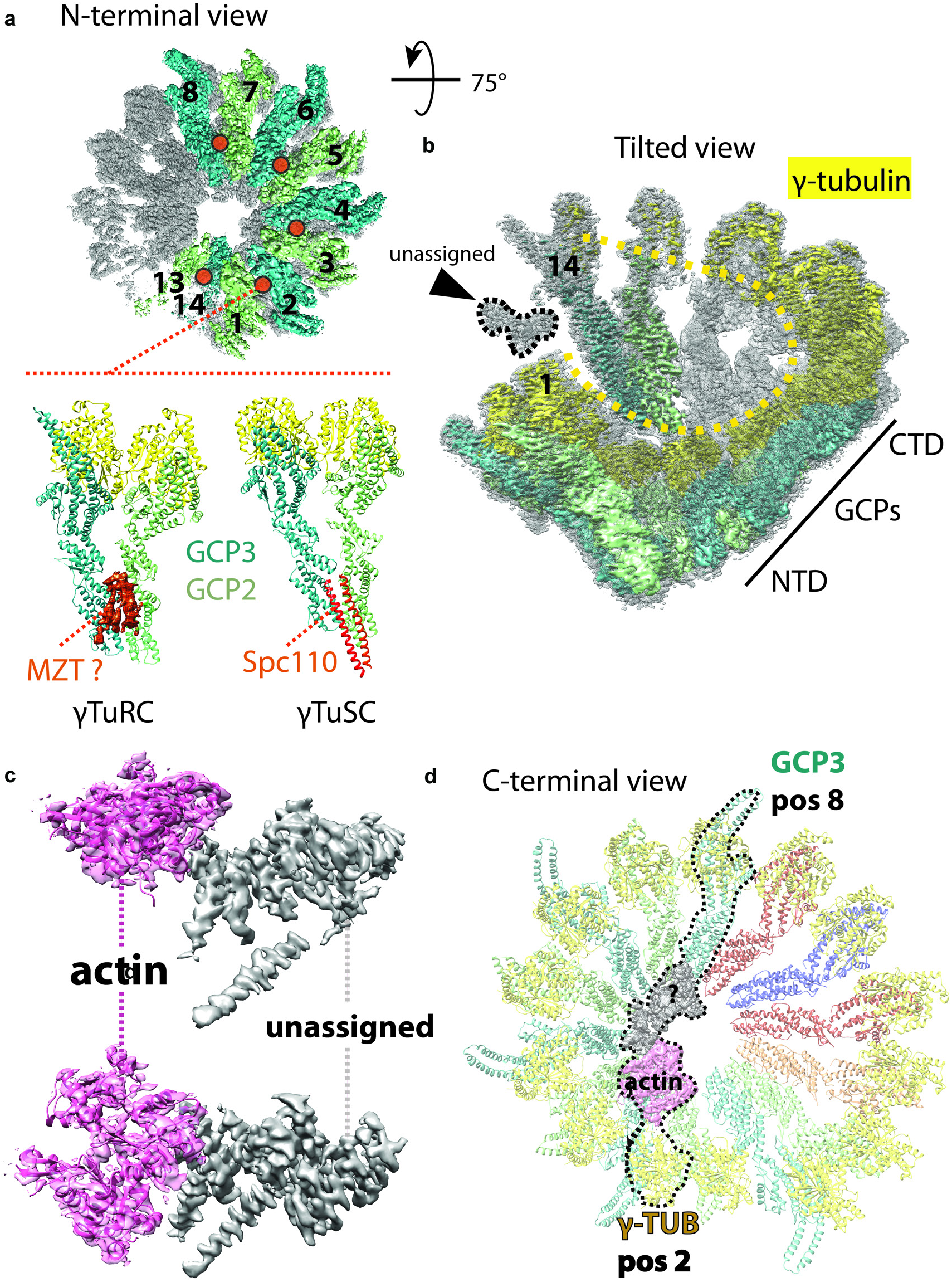
Analysis of the unassigned cryo-EM density in the γTuRC complex. **(a)** A characteristic V-shaped *α*-helical feature appears to seal off the interface of GCP2 and GCP3, lining the outer perimeter of the GCP spiral. This feature occupies the same position observed for SCP110 in γTuSC. We propose that this density could be MZT1 or MZT2, which were both found in our γTuRC preparation. **(b)** Unassigned density can be observed departing from the C-terminal end of GCP3 in position 14 (marked with a black arrowhead). **(c)** Additional density is found in the lumen of the γTuRC spiral. Part of this density can be assigned to actin (magenta), which was found to be co-purified in our preparation. Additional unassigned density shown in grey contains three recognisable alpha helical bundles. **(d)** The luminal density bridges γ-tubulin in position 2 and GCP3 in position 8.

Additional, prominent density could also be observed within the lumen of the γTuRC cone. In line with previous reports on γTuRC purification (Choi et al., 2010; Oegema et al., 1999), both actin featured in our list of proteins co-purified in our preparation. We therefore attempted docking one actin protomer (PDB entry 2HF3 (Rould et al., 2006)) into the luminal density (Fig. 7c). Our attempt resulted in an excellent fit, leading us to establish that actin is a structural component of the γTuRC complex, contacting the γ-tubulin subunit in position 2. Actin also interacts with one unassigned luminal feature, formed by a helical repeat module, which straddles across the central pore of the helical assembly, bridging between γ-tubulin in position 2 with GCP3 in position 8 (Fig. 7d). Thus, a combination of AHF proteins lining the outer perimeter of GCP subunits in position 1-8 appear to seal off GCP2-3 interfaces, and function together with actin and associated luminal factors poised to rigidify a large portion of the left-handed spiral of γTuRC. We note that γTuRC protomers in position 1-8 match the GCP2-3 configuration in the active (“closed”) configuration of yeast γTuSC (Kollman et al., 2015). Conversely, subunits in positions 9-14, which notably lack any bridging element within the spiral lumen, display a configuration more akin to the inactive (“open”) form of yeast γTuSC (Kollman et al., 2015). Coherently, subunits in positions 1-8 accurately match the geometry of a 13-subunit microtubule protofilament, and could hence sustain nucleation (Fig. S8). The observation that γTuRC subunits in position 9-14 markedly diverge from microtubule geometry could justify our observation that γTuRC enhanced microtubule nucleation is kinetically inhibited.

## DISCUSSION

Using TIRF microscopy, we imaged the nucleation of individual microtubules by surface-immobilized human γTuRC. Microtubules were stably capped by γTuRC, not displaying any minus end dynamics. γTuRC increased the nucleation efficiency compared to microtubule nucleation in solution, but nucleation was still kinetically inhibited. Microtubule formation on the γTuRC template was mechanistically very different from microtubule plus end elongation. Determining the structure of the human γTuRC complex at 4 Å resolution revealed an asymmetric conformation with half of the complex in a compact configuration (closed) and half containing loosely interacting protomers (open). This structure is distinctly different from the structure of budding yeast γTuRC (Kollman et al., 2015), particularly where the GCP4, 5 and 6 subunits are located that are absent in yeast γTuRC.

Microtubule nucleation in solution has been described as a highly cooperative process. Many tubulins need to come together to form a first minimal stable assembly (Voter and Erickson, 1984). Although γTuRC is thought to nucleate microtubules by providing a template that mimics the microtubule structure, also nucleation from this template is kinetically inhibited and highly cooperative. Our estimate of at least ∼ 7 tubulins needing to come together before stable microtubule outgrowth from the γTuRC complex can occur (Fig. 3d) is at the lower end of the reported range of 6-15 tubulins for such a critical nucleus required for spontaneous nucleation in solution (Flyvbjerg and Jobs, 1997; Fygenson et al., 1995; Voter and Erickson, 1984), supporting the notion that the template facilitates nucleation.

However, microtubule outgrowth from the γTuRC template is considerably more difficult than elongation from a pre-existing growing microtubule plus end. Only a fraction of γTuRCs nucleated within 20 min in our assay. We did not observe any indication for a permanently inactive γTuRC population, because the probability of stochastic nucleation was constant over time, indicating that the majority of theγTuRC molecules may be in a inhibited conformation.

Our γTuRC structure supports this notion. Only half of the human γTuRC complex exists in a closed configuration with four GCP2/3 heterodimers bound to 8 γ-tubulins being bridged together by stabilising factors including one actin monomer in the lumen of the cone-shaped complex. Conversely, the other half of the complex where GCP4/5/6 subunits are located, together with one additional GCP2/3 dimer at the very top of the helical arrangement of GCP dimers, lack a bridging luminal elements and only loosely interact with one another. This open configuration causes a γ-tubulin arrangement that deviates from the microtubule geometry and that is expected for an active γTuRC state (Movie 8). This asymmetric structure indicates that the purified human γTuRC is not in a fully active conformation. Either microtubule assembly on the γTuRC surface may induce γTuRC closure and/or additional regulatory binding factors may induce a completely closed γTuRC geometry required for efficient nucleation, suggesting a mechanism for the regulation of γTuRC activity.

In favour of the scenario of microtubule assembly-induced γTuRC closure is our observation that proteins which stabilize growing microtubule ends by different means and which were shown to stimulate microtubule nucleation *in vitro*, also stimulate γTuRC-mediated microtubule nucleation. The microtubule polymerase chTOG may do so by accelerating and thereby stabilising nascent microtubule growth on γTuRC by its polymerase activity (Brouhard et al., 2008; Roostalu et al., 2015), and TPX2 may do so by its catastrophe suppressing activity (Roostalu et al., 2015; Wieczorek et al., 2015). chTOG and TPX2 were also reported to promote microtubule outgrowth from stabilised microtubule ‘seeds’, suggesting similarities between outgrowth from such seeds and templating a microtubule by γTuRC (Wieczorek et al., 2015). Interestingly, both chTOG and TPX2 have also been reported to directly interact with γTuRC, which may increase the efficiency of their action (Alfaro-Aco et al., 2017; Thawani et al., 2018). Moreover, TPX2 is known to interact additionally with the Augmin/HAUS complex that promotes branched microtubule nucleation from pre-existing microtubules in mitotic or meiotic spindles during cell division (Alfaro-Aco and Petry, 2017).

Our structure suggests a potential mechanism for the activation of human γTuRC by complex closure. In our structure, all GCP2/3 interfaces in the closed part of the complex are sealed off by inter-protomer elements, in a position reminiscent of yeast γTuSC specific Spc110, and which we tentatively assign to MZT factors present in our preparation. As for Scp110, we propose that MZT factors are likely required to stabilise the GCP2/3 spiral formation at the constricted N-terminal dimerization core. At least 5 additional binding sites in the open part of the complex remain available at inter-protomer interfaces along the perimeter of the constricted spiral base, which could be engaged by additional nucleation-activation factors.

These elements in the γTuRC structure suggest an activation mechanism for human γTuRC that differs from the activation mechanism for budding yeast γTuRC. This difference is likely a consequence of different GCP subunit compositions of the two complexes. Yeast γTuRC consists of a helical arrangement of 7 identical γTuSCs (2 γ-tubulins and one GCP2 and 3 each) and displays gaps between every second γ-tubulin (within a γTuSC) that create a mismatch with the microtubule geometry (Kollman et al., 2015). Consequently, in addition to regulation at the level of complex assembly by recruitment factors to the spindle pole body, yeast γTuRC can be further activated by a structural change closing the gaps between the γ-tubulins, resulting in a γ-tubulin arrangement that matches the 13-mer protofilament geometry of the microtubule (Kollman et al., 2015). In contrast, the human γTuRC structure departs from the microtubule geometry where GCP4, 5 and 6 are located, resulting in an entire half of the complex being in an open conformation. It is tempting to speculate that other proteins that may line the outer perimeter of the open part of γTuRC (or its inner lumen) may induce a conformational change, closing the second half of the complex and thereby producing a completely closed conformation in which the γ-tubulin configuration closely matches the geometry of the microtubule lattice. Such reconfiguration may consequently reduce the kinetic barrier for templated microtubule nucleation.

We note that our cryo-EM reconstruction results closely match those recently reported in a bioRxiv submission (Wieczorek et al., 2019), yet the interpretation of the map presents some differences. Although the AHF elements on the outer perimeter of the GCP cone appear very similar in both cryo-EM maps, they were assigned to γTuNA (a CDK5Rap2 fragment) in the bioRxiv study, as this factor was used for affinity purification of the γTuRC complex. Our purification strategy did not include γTuNA and this factor was not detected in our γTuRC preparation, leading us to assign this cryo-EM feature to MZT proteins, which were present in our preparation. Since Spc110 binds to the same position in budding yeast γTuRC, this may indicate a promiscuous binding site in γTuRC that may be available to a range of regulators. Another interesting difference with the bioRxiv study is observed at the GCP6/GCP2 inter-protomer interface, which contains a structured element assigned to a GCP6 insertion in the Wieczoreck et al. study, but is notably absent in our cryo-EM map. We speculate that this element might be a stabilizing factor, which was not co-purified in our preparation. Alternatively, it might be indeed GCP6, captured in a different configuration in the two structures.

A major open question for the future will be to understand how the various proteins involved in controlling the efficiency of microtubule nucleation in cells control the conformation of human γTuRC. Our real-time *in vitro* nucleation assay in combination with structural investigations will be essential to shed light on the detailed mechanism of spatio-temporal γTuRC activation by an open-to-closed transition.

## Supporting information

Movie1_Consolati

Movie2_Consolati

Movie3_Consolati

Movie4_Consolati

Movie5_Consolati

Movie6_Consolati

Movie7_Consolati

Movie8_Consolati

SupplementaryMaterial_Consolati

## ACKNOWLEDGEMENTS

We thank Andrea Nans at the Structural Biology Science Technology Platform (STP) for support on the Titan Krios, Raffa Carzaniga at the Electron Microscopy STP of the Francis Crick Institute for support on the Tecnai G2 Spirit electron microscope, Nicholas I. Cade for fluorescence microscopy support, and Andrew Purkiss and Phil Walker for computational support. We thank the Cell Sevices, the Flow Cytometry and the Mass Spectrometry Proteomics STPs of the Francis Crick Institute for producing large cell cultures, and for cell sorting and protein identification support. This work was supported by the Francis Crick Institute, which receives its core funding from Cancer Research UK (FC001163, FC0010065), the UK Medical Research Council (FC001163, FC0010065), and the Wellcome Trust (FC001163, FC0010065) to T.S. and A.C. J.R. was supported by a Sir Henry Wellcome Postdoctoral Fellowship (100145/Z/12/Z) and T.S. acknowledges support from the European Research Council (Advanced Grant, project 323042). A.C. receives funding from the European Research Council (ERC) under the European Union’s Horizon 2020 research and innovation programme (grant agreement No 820102). T.C., J.W.M. and T.S. acknowledge also the support of the Spanish Ministry of Economy, Industry and Competitiveness to the CRG-EMBL partnership, the Centro de Excelencia Severo Ochoa and the CERCA Programme/Generalitat de Catalunya.

## AUTHOR CONTRIBUTIONS

Conceptualization, T.C., J.L., J.R., A.C. and T.S.; Methodology, T.C., J.L., J.R., J.A. and W.M.L.; Validation, T.C. and J.L.; Formal Analysis, T.C., J.L., F.M. and A.C.; Investigation, T.C., J.L., J.A., W.M.L., J.G.; Resources, J.R. and J.G., Original Draft, T.C., A.C. and T.S.; Review & Editing, T.C., J.A., W.M.L., A.C., and T.S.; Visualization, T.C., J.L., F.M., A.C.; Supervision, J.R., A.C. and T.S., Project Administration, T.C., J.L., A.C. and T.S.; Funding Acquisition, A.C. and T.S.

## EXPERIMENTAL PROCEDURES

### Lentivirus expression constructs and molecular biology

To generate a fluorescently-tagged and biotinylatable human γTuRC, the coding region for full-length human GCP2 (amino acids 1-902) was amplified by PCR using its cDNA as template (NM_001256617.1, Origene). The mTagBFP (blue fluorescent protein, Evrogen) coding sequence was also amplified by PCR. Both PCR-amplified sequences were cloned into a pLVX-Puro vector (Clonetech) using Gibson assembly (In-Fusion cloning, Takara), to form GCP2_G_5_A_TEV_G_5_A_mTagBFP_G_5_A_BAP, an expression construct for GCP2 which is C-terminally tagged with mTagBFP and biotin acceptor peptide (BAP: GLNDIFEAQKIEWHE), both separated from GCP2 by a TEV protease cleavage site. Glycine linkers (G_5_A) were placed between sequences. To facilitate the *in vivo* biotinylation of tagged γTuRC *E. coli* biotin ligase BirA was cloned into a pLVX-IRES-Hyg vector (Clonetech) using Gibson assembly to form HA_ G_5_A_BirA; an expression construct of BirA with an HA-tag added to the BirA N-terminus, separated by a G_5_A-linker. Primers used for cloning are listed in Table S1.

### Antibodies

Commercial and custom-made antibodies were used for the characterization of purified γTuRC by western blotting (Table S2). Custom-made antibodies were raised against His_6_-tagged proteins expressed and purified from *E. coli*. Specific antibodies were affinity purified by standard methods using MBP-tagged proteins expressed and purified from *E. coli* and coupled to CNBr-beads (GE Healthcare). The specificity of custom-made antibodies was confirmed by western blotting against human cell lysate after RNAi depletion of target proteins as described previously (Cota et al., 2017). For detection of biotinylated proteins by western blot, peroxidase coupled streptavidin (streptavidin-HRP, Thermo Fisher) was used.

### Cell culture and cell line development

HeLa-Kyoto cells (RRID:CVCL_1922) were cultured at 37°C (10% CO_2_) in Dulbecco’s Modified Eagle Medium (DMEM) supplemented with 10% fetal bovine serum, 50 U mL^-1^ penicillin and 50 µg mL^-1^ streptomycin. To generate HeLa-Kyoto cells stably expressing biotinylated mTagBFP-tagged GCP2, cells were co-transduced with GCP2 and BirA lentivirus (Abella et al., 2016) followed by hygromycin and puromycin selection. Resistant cells expressing mTagBFP were sorted by FACS (fluorescent assisted cell sorter) and cultured independently in 96 well plates. The isolated single-cell colonies were screened for HA-BirA expression by immunofluorescence staining (primary antibody: mouse anti-HA (F-7, Santa Cruz Biotechnology); secondary antibody: goat anti-mouse-FITC (Sigma)) and then using high throughput imaging (High throughput screening facility, Francis Crick Institute). The localisation of GCP2-mTagBFP-BAP was confirmed by live-cell fluorescence imaging using a spinning disc confocal microscope based on a NikonTI-E frame with a 100x 1.49 N.A. Nikon objective lens (Cairn Research, Faversham, UK). mTagBFP expressing colonies were further tested by western blotting to confirm the expression of GCP2-mTagBFP-BAP and HA-BirA.

When producing large cell cultures for purification, three days before harvesting cells (using trypsination), D-biotin (Sigma Aldrich) was added to a final concentration of 50 μM. Cell pellets were stored at −80°C until further use.

### Purification of human γTuRC

Cells were resuspended in lysis buffer (50 mM HEPES, 150 mM KCl, 5 mM MgCl_2_, 1 mM EGTA, 1 mM DTT, 0.1 mM GTP, pH 7.4) containing protease inhibitors (complete EDTA-free protease inhibitor mix, Roche) and DNAse I (10 μg ml^-1^, Sigma-Aldrich). Resuspended cells were lysed using a polytron tissue dispenser (3×90 s at 6.6×10^3^ rpm) and lysate was clarified twice by centrifugation (17,000xg, 15 min, 4°C). Clarified lysate was filtered through three sets of filters with decreasing pore size: 1.2 µm (GE Healthcare), 0.8 µm (GE-Healthcare) and 0.45 µm (Millipore). The lysate was buffer exchanged into storage buffer (lysis buffer containing 0.02% (vol./vol.) Brij-35) over HiPrep 26/10 desalting columns to remove D-biotin from the lysate. Protein-containing fractions were pooled, supplemented with protease inhibitors and loaded onto a 1 mL HiTrap SP Sepharose FF column connected in tandem with 1 mL streptavidin mutein matrix beads (Sigma Aldrich) packed into a Tricorn 5/50 column. The streptavidin mutein matrix column was washed with 30 mL storage buffer, 30 mL wash buffer (lysis buffer containing 200 mM KCl and 0.2% (vol./vol.) Brij-35 and) and 30 mL storage buffer. Proteins were eluted with storage buffer supplemented with 5 mM D-biotin. The buffer was then exchanged back into storage buffer using a HiTrap Desalting column. Protein-containing fractions were pooled and concentrated using Amicon centrifugal units (MWCO 30,000, Millipore), centrifuged (17,000xg, 10 min, 4°C) and separated by size exclusion chromatography using a Superose 6 10/300 GL column. γTuRC peak fractions were pooled, concentrated, ultracentrifuged (278,088.3xg, 10 min, 4°C), snap frozen and stored in liquid nitrogen. From 120 g of cell pellet typically ∼85 μg of tagged γTuRC were purified.

### Purification of human chTOG-mGFP, mGFP-TPX2 and mGFP-EB3

GFP-tagged microtubule binders were purified as described (Jha et al., 2017; Roostalu et al., 2015).

### Tubulin purification and labeling

Porcine brain tubulin was purified and covalently labelled with NHS-biotin (Thermo Fisher) or NHS-CF640R (Sigma-Aldrich) as described (Castoldi and Popov, 2003; Hyman et al., 1991).

### LC-MS/MS analysis of fluorescently tagged γTuRC

Purified γTuRC was separated by SDS-PAGE and stained using InstantBlue (Expedeon). Protein bands were excised from the gel and analysed by the Francis Crick Institute Proteomics facility. Briefly, Tryptic peptides were analysed using a Q Exactive orbitrap mass spectrometer coupled to an Ultimate 3000 HPLC equipped with an EasySpray nano-source (Thermo Fisher Scientific). A one hour method of MS1 orbitrap (60k resolution) followed by top 10 HCD MS2 (35k resolution) produced raw data files. Raw files were analysed in MaxQuant (v1.6.0.13) against the SwissProt Homo sapiens protein database (downloaded June 2019) using the iBAQ algorithm. The canonical GCP2 sequence was replaced with the construct sequence (GCP2-5xGly-TEV-5xGly-mBFP-5xGly-AviTag). Variable modifications of methionine oxidation and protein N-terminal acetylation along with a fixed modification of cysteine carbamidomethylation were selected. The proteingroups.txt file was imported in Perseus (v1.4.0.2) for data analysis. Potential contaminants, reverse sequences and proteins identified by site were removed. iBAQ intensities were log2 transformed.

### TIRF microscopy-based microtubule nucleation assay

The γTuRC-mediated microtubule nucleation assay was modified from a previous surface nucleation assay without γTuRC (Roostalu et al., 2015). Flow chambers were assembled from biotin-polyethylene glycol (PEG)-functionalized glass and poly(L-lysine)-PEG (SuSoS)-passivated counter glass as described previously (Bieling et al., 2010).

A flow chamber was incubated for 10 min with 5% Pluronic F-127 in MilliQ water (Sigma Aldrich), washed with assay buffer (AB: 80 mM PIPES, 60 mM KCl, 1 mM EGTA, 1 mM MgCl_2_, 1 mM GTP, 5 mM 2-ME, 0.15% (w/vol.) methylcellulose (4,000 cP, Sigma-Aldrich) 1% (w/vol.) glucose, 0.02% (vol./vol.) Brij-35)) supplemented with 50 μg mL^-1^ κ-casein (Sigma-Aldrich), followed by a 3-min incubation with the same buffer additionally containing 50 μg mL^-1^ of NeutrAvidin (Life Technologies). The chamber was subsequently washed with γTuRC storage buffer and incubated for 5 min with prediluted γTuRC in γTuRC storage buffer to the concentration indicated for each experiment. Unbound γTuRC was removed by washing the flow cell with AB. Then the final assay mix was passed through, the chamber was sealed with vacuum grease (Beckman) and placed onto the microscope.

Final assay mix: AB supplemented with oxygen scavengers (160 μg mL^-1^ catalase (Sigma-Aldrich), 680 μg mL^-1^ glucose oxidase (Serva)) diluted in BRB80 (80mM PIPES, 1mM EGTA, 1mM MgCl_2_), 1 mg ml^-1^ bovine serum albumin (Sigma-Aldrich) in BRB80, varying concentrations of tubulin (containing 4.8% CF640R-labelled tubulin). For experiments with microtubule binders 2.9% (vol./vol.) of either chTOG-mGFP, mGFP-TPX2 or mGFP-EB3 was added at different concentrations. chTOG-mGFP and mGFP-TPX2 concentrations were altered by predilution in their storage buffers (Jha et al., 2017; Roostalu et al., 2015). mGFP-EB3 was diluted in BRB80. The final assay mix containing chTOG-GFP was ultracentrifuged (278,088.03xg, 10 min, 4°C) before flowing the mix into the chamber. To keep the buffer composition of the final assay mix unchanged within a set of experiments and to allow for direct comparisons between experiments, the overall BRB80 and storage buffer content was kept constant within one set of experiments.

All experiments were performed using a total internal reflection fluorescence (TIRF) microscope (Cairn Research, Faversham, UK) (Hannabuss et al., 2019). Experiments were imaged 2 min after placing the chamber on the microscope. The temperature was kept at 33±1°C. Two-and three-colour time-lapse imaging for γTuRC nucleation assays and seed assays were performed at 1 frame/5 s with a 1-s exposure time using a 60x 1.49 NA Nikon objective lens. For single molecule γTuRC assays shown in Fig. 2f, images were acquired at 1 frame/1.8 s with a 500-ms exposure time using a 100x 1.49 N.A. Nikon objective lens. CF640R-tubulin (640 nm excitation) and mGFP-tagged proteins (488 nm excitation) were imaged simultaneously. γTuRC-mTagBFP-BAP (405 nm excitation) was imaged every 10 frames for single molecule γTuRC assays and once at the beginning and at the end of the movie for γTuRC nucleation assays.

### TIRF microscopy image processing

The Fiji package of ImageJ was used to generate kymographs (space-time plots) and to merge image sequences from different channels. For multi-colour imaging, image alignment was performed using Matlab as described (Maurer et al., 2014). Background was subtracted using the background subtraction tool of Fiji (‘rolling ball’ method). For movies from single molecule γTuRC assays shown in Fig. 2f γTuRC-mTagBFP-AviTag images were merged using the ‘grouped Z project’ function in Fiji. To subtract camera noise an empty flow chamber was imaged using the same imaging conditions. The background image was generated as described above and subtracted from the γTuRC-mTagBFP-AviTag image, which was then used to merge with images of CF640R-tubulin.

**Microtubule growth speeds** were measured directly from kymographs using the ‘Resclice function’ in Fiji. Lines were drawn manually along growing plus- and minus-ends. Growth speeds were calculated from the slope of the line.

### Microtubule nucleation rate analysis

For each nucleation assay, microtubules were counted manually at 10 different time points either until the end of the movie or until individual microtubule nucleation events could no longer be identified due to overcrowding. The total number of nucleated microtubules in a field of view at a given time point was obtained by counting the newly nucleated microtubules and adding it to the number obtained at the previous analysed time point. Microtubule numbers were tracked using the ‘Point tool’ together with the ‘ROI manager tool’ in Fiji. For the quantification of γTuRC-mediated microtubule nucleation rates, only microtubules were counted that started nucleating from the surface and that stayed surface-attached. Microtubule nucleation rates represent the slope of the linear regression for each condition and are given in number per nucleated microtubules per field of view and per time.

### Nucleation and growth statistics

All error bars represent the standard error of mean (s.e.m.) or standard deviation (s.d.) as indicated in each Figure. Linear regression and curve fitting were performed using Prism software (GraphPad).

### Negative stain grid preparation and data collection

A 4-µl droplet of purified γTuRC diluted in γTuRC storage buffer was applied to a freshly glow-discharged carbon-coated grid (C300Cu100, EM Resolution) and incubated for 2 min. The grid was stained with consecutive applications onto three 50-µl droplets of 2% uranyl acetate solution for 30 s each. The grid was then blotted dry and stored until imaged on a 120 keV G2 Spirit transmission electron microscope (FEI) equipped with a 2k×2k Ultrascan-1000 camera (Gatan). The Micrographs were collected using a nominal magnification of 30,000x, resulting in a pixel size of 3.45 Å at the specimen level.

### Cryo grid preparation and data collection

Freeze-thawed purified γTuRC was briefly spun to remove aggregates. Lacey grids (400 mesh) with a layer of ultra-thin carbon (Agar Scientific) were glow-discharged at 45 mA for 1 min using a K100X Glow Discharge Unit (EMS). A 4 µl-droplet was then applied directly onto the carbon-side of the grid loaded into the humidity chamber of a Vitrobot Mark IV (Thermo Fisher) set to room temperature and 90% humidity. After an incubation time of 60 seconds, the grid was blotted for 3.0 s and plunged into liquid ethane. The ice quality was assessed on a 200 kV Talos Arctica (Thermo Fisher) and a small dataset was collected to evaluate the sample quality.

The highest-quality grid was imaged using a 300kV Titan Krios electron microscope (Thermo Fisher) using a GIF Quantum energy filter (Gatan) and a K2 Summit direct detector (Gatan), operated in counting mode. A total of 2,4000 movies were collected over two sessions at a pixel size of 1.08 Å/px with a total dose of ∼50 e−/A2 and a defocus range of −1.0 − −3.5 µm.

### Negative stain electron microscopy image processing

The particles were picked using e2boxer.py of the EMAN2 v2.07 software package (Tang et al., 2007), using the semi-automated (swarm) option. Box files were then imported in the RELION-3.0 (Zivanov et al., 2018), which was used for all downstream image processing steps were performed. Contrast transfer function parameters were determined using Gctf v.1.18 (Zhang, 2016), and extracted particles were subjected to two-dimensional classification.

### Cryo-EM image processing

To correct for beam-induced movements all movie frames were aligned using dose-weighted averaging in MotionCor2(Zheng et al., 2017). CTF parameters were estimated using non-dose-weighted micrographs generated by Gctf v.1.18 (Zhang, 2016). Automated particle-picking was performed using crYOLO of the SPHIRE software package (Wagner et al., 2019). Box files were imported in RELION-3 (Zivanov et al., 2018) and a total of ∼ 1.1 million particles were initially binned by a factor of four and extracted from dose-weighted micrographs with a box size of 128 pixels. After several rounds of two-dimensional classification, a total of 522,496 high-resolution particles were selected, which evidently contained high-resolution information. Unbinned particles were re-extracted, using a 515-pixel box size. These particles were used to generate three reference free *ab initio* models using cryoSPARC v2 (Punjani et al., 2017). The best model, which resulted from 229,744 particles, was imported in RELION-3, filtered to 60 Å and used as a starting reference for 3D classification of the 522,496 high-resolution particles. The combination of all particles yielded in the highest resolution class, which was subsequently subjected to one initial 3D refinement, followed by three rounds of CTF refinement and one Bayesian particle polishing step. Polished particles were subjected to one final round of CTF refinement, 3D refinement and post processing, yielding in a final 3D structure with an overall resolution of 3.98 Å.

### Generation of an atomic model

The crystal structure of human GCP4 (PDB entry 3RIP) (Guillet et al., 2011) was separated in three distinct domains and used for docking into the cryo-EM map, using the Fit in map option in UCSF Chimera (Pettersen et al., 2004). Highest correlation GCP subunits were assigned to GCP4, while GP3 was recognised because of a characteristic helical extension in the C-terminal γ-tubulin interacting domain, first modelled using iTasser (Roy et al., 2010), adjusted manually in Coot (Emsley et al., 2010) and refined using Phenix (Afonine et al., 2018) and Namdinator (Kidmose et al.).

---

**Figure S1.**
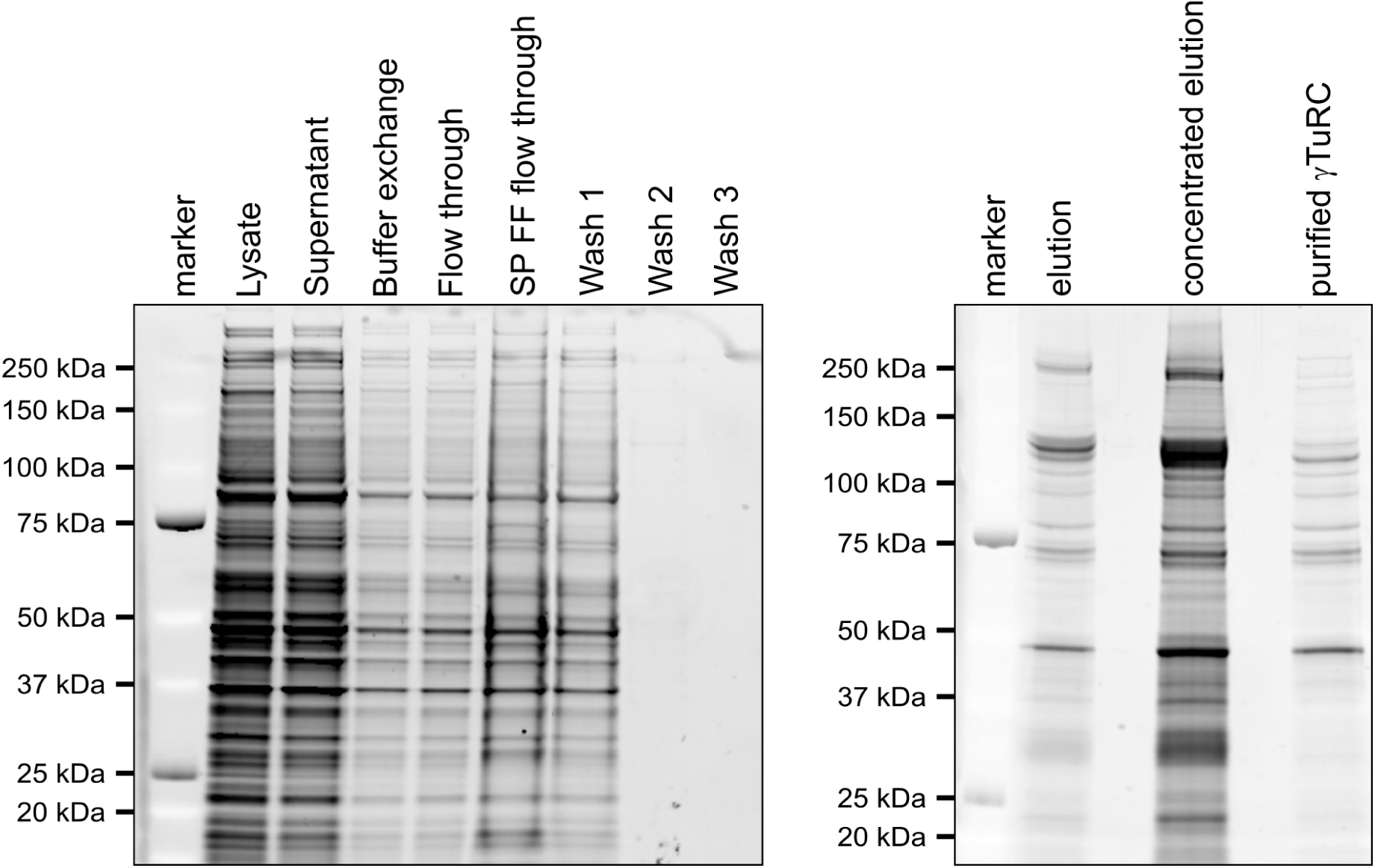
Purification of tagged human γTuRC. Sypro-Ruby stained SDS-PAGE showing the purification of γTuRC-mBFP-BAP purification from human cells.

**Figure S2.**
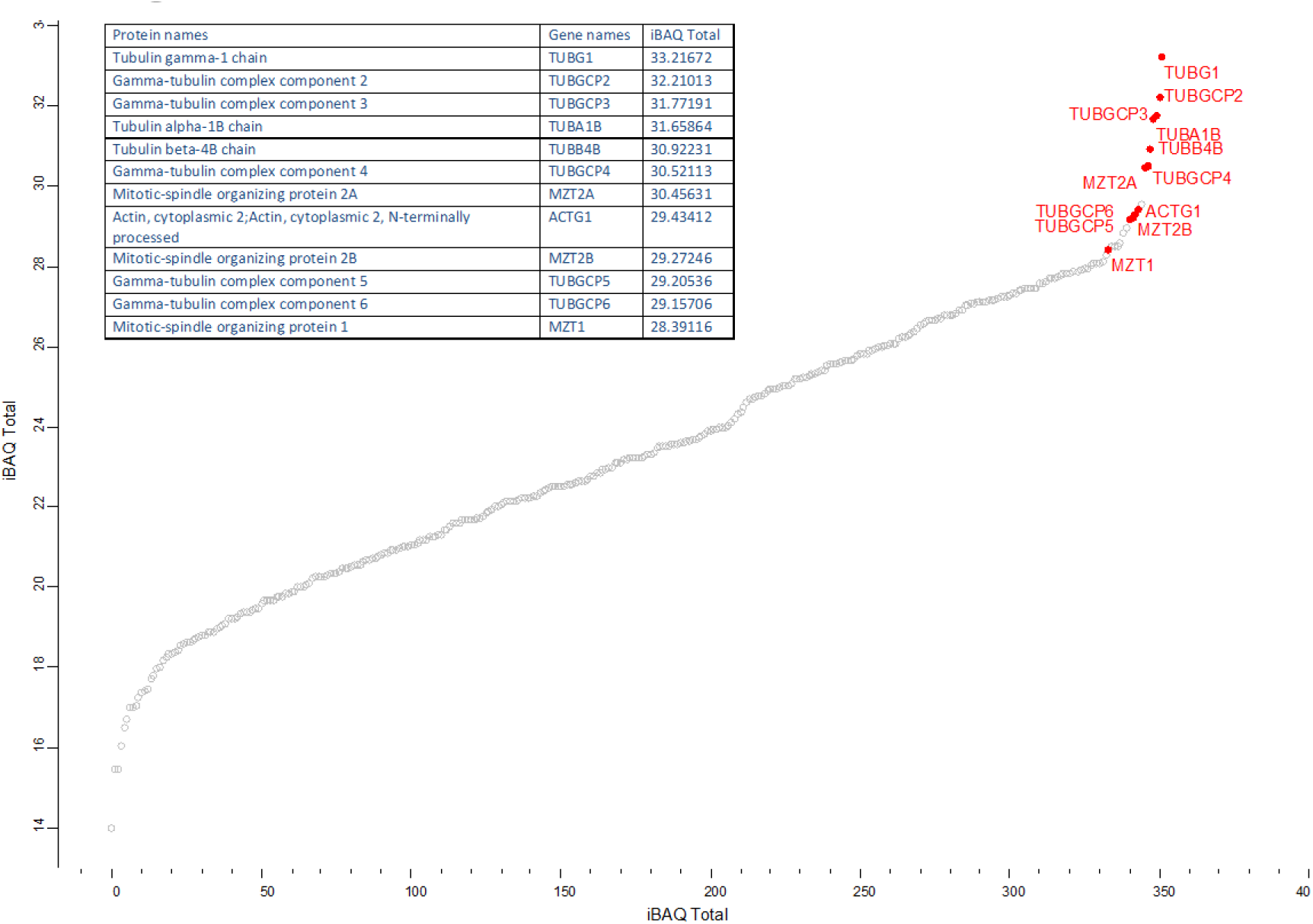
Scatter plot representing the protein density by iBAQ intensity.

**Figure S3.**
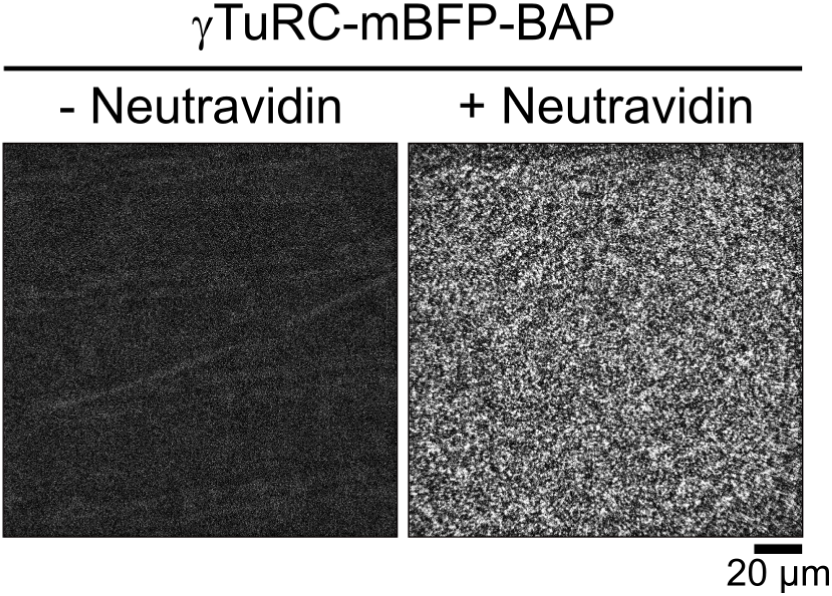
Specificity of γTuRC immobilization on functionalized glass surfaces. Biotinylated fluorescent γTuRC was bound to functionalized glass surfaces pre-incubated with NeutrAvidin or in the absence of NeutrAvidin as indicated. Representative TIRFM images of γTuRC surface density (mBFP fluorescence) are shown. Scale bar as indicated.

**Figure S4.**
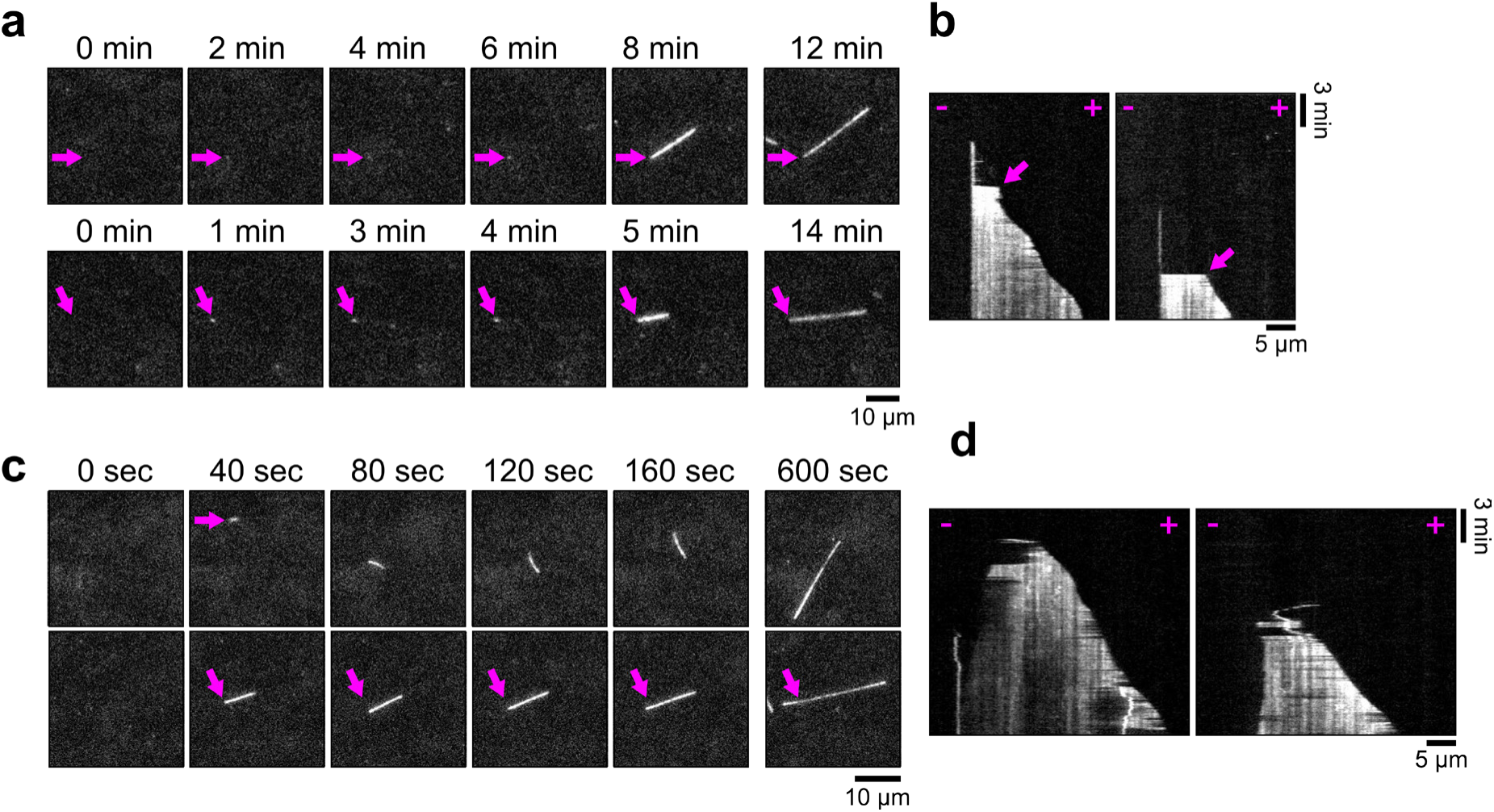
Examples of ‘falling’ γTuRC-nucleated microtubules and ‘landing’ spontaneously nucleated microtubules in solution. **(a)** Representative time series of two γTuRC-nucleated microtubules that first grow out of the TIRF field and then ‘fall’ onto the surface (15 μM CF640R-tubulin, 93 pM γTuRC). ‘Falling’ microtubules only grow from one end while the other end is stably anchored to the surface indicating that these microtubules are initiated by surface immobilized γTuRC. Purple arrows indicate the stably anchored microtubule ends. **(b)** Corresponding TIRFM kymographs. Purple arrow marks ‘falling’ event. **(c)** Representative time series of two spontaneously nucleated microtubules, ‘landing’ from solution (15 μM CF640R-tubulin, 47 pM γTuRC). Spontaneously nucleated microtubules are distinguishable from γTuRC-nucleated microtubules because they are not anchored to the surface (top row, purple arrow mark first appearance of spontaneously nucleated microtubule) and grow with both ends (bottom row, purple arrows mark slow growing minus-end). **(d)** Corresponding TIRFM kymographs. Scale bars as indicated. t=0 is 2 min after placing the sample at 33 °C.

**Figure S5.**
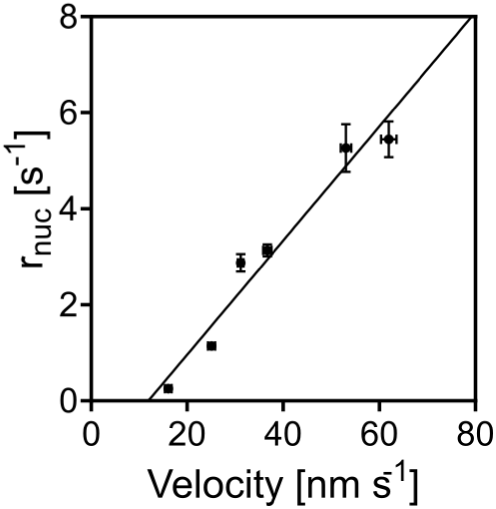
Linear relationship between γTuRC-mediated microtubule nucleation and chTOG-accelerated microtubule plus-end growth. Plot of the mean nucleation rate (r_nuc_) against the mean microtubule plus-end growth speed at different mGFP-chTOG concentrations. Data were pooled from at least 3 data sets. Line represents the linear regression. Error bars are s.e.m.

**Figure S6.**
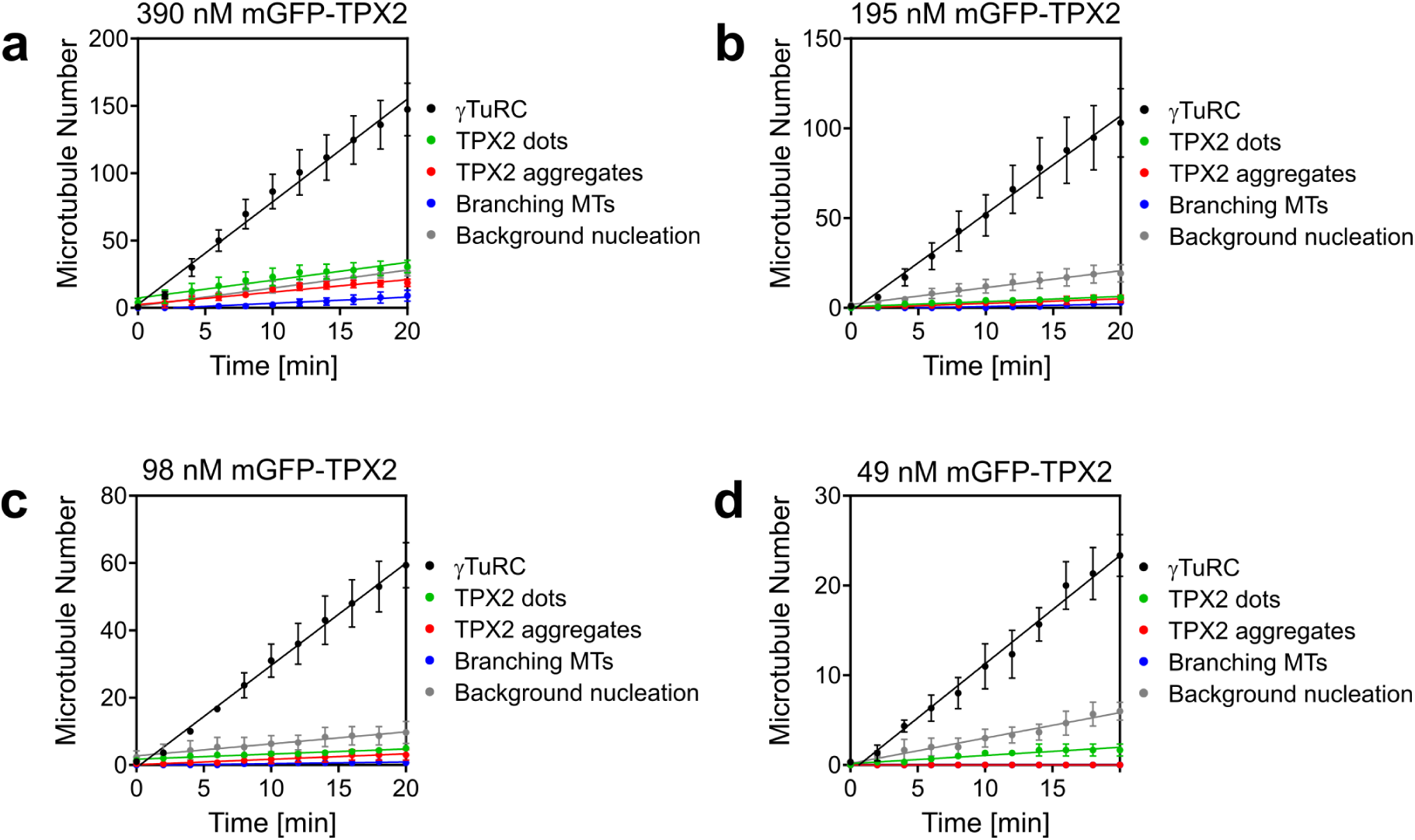
Microtubules nucleated in assays performed in the presence of mGFP-TPX2. Plots showing linearly increasing nucleated microtubule numbers over time at different mGFP-TPX2 concentrations, as indicated. For all conditions, the majority of microtubules are nucleated by γTuRC (black line). Non-γTuRC nucleated microtubules were categorized as microtubules growing from ‘dots’ of mGFP-TPX2 (green line), microtubule bundles growing out from large TPX2 aggregates (red line), microtubules branching from existing microtubules with TPX2 bound (blue line) and spontaneously nucleated microtubules (grey line).

**Figure S7.**
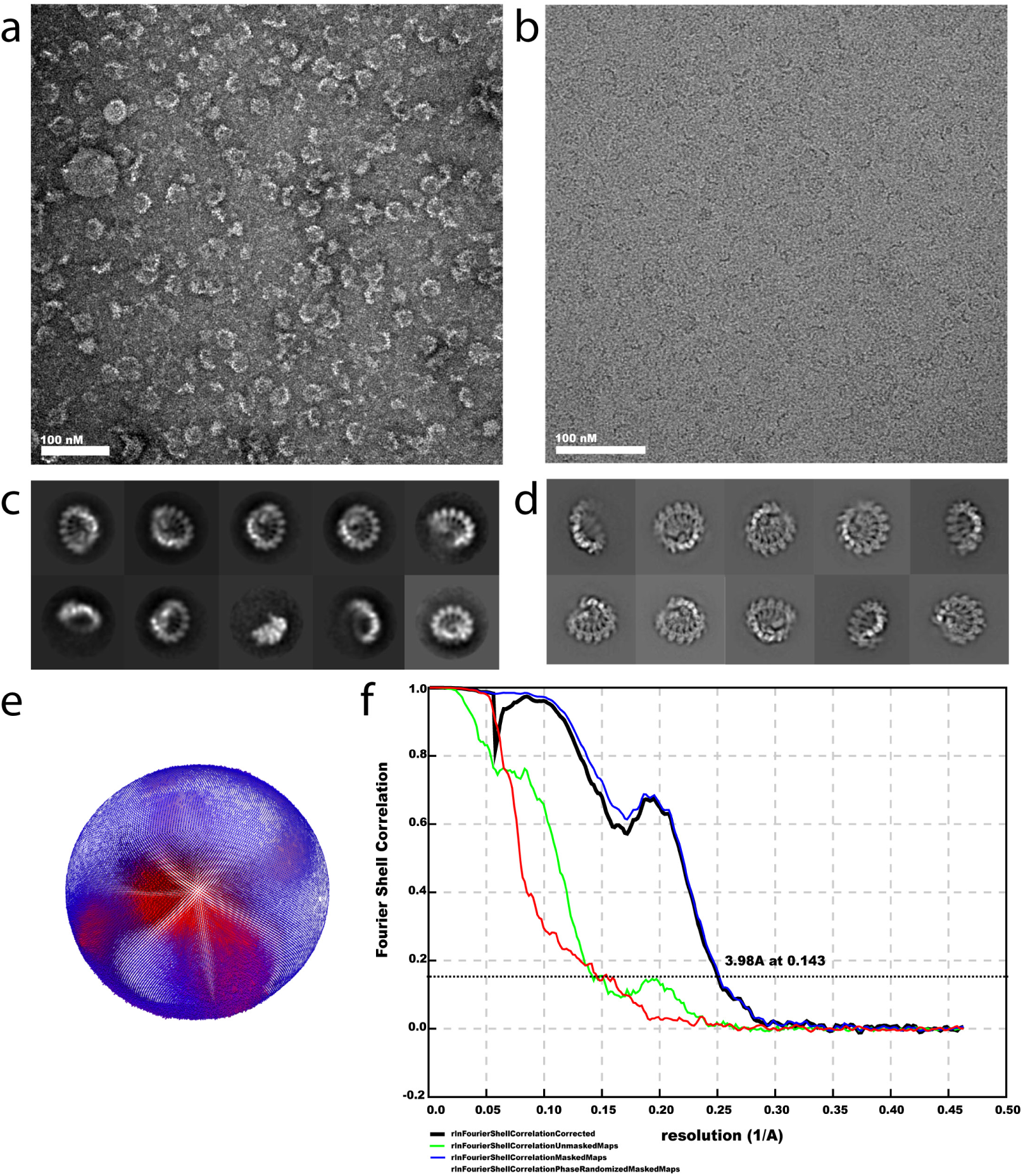
Electron microscopy image acquisition and processing. **(a)** Micrograph of a negatively stained sample. **(b)** Cryo-electron micrograph of a frozen-hydrated sample. **(c)** 2D class averages deriving from negative stain images. **(d)** 2D class averages deriving from cryo-EM images. **(e)** Angular distribution of particles used in the final cryo-EM reconstruction. **(f)** Fourier shell correlation indicating a resolution of 3.98 Å according to the 0.143 criterion.

**Figure S8.**
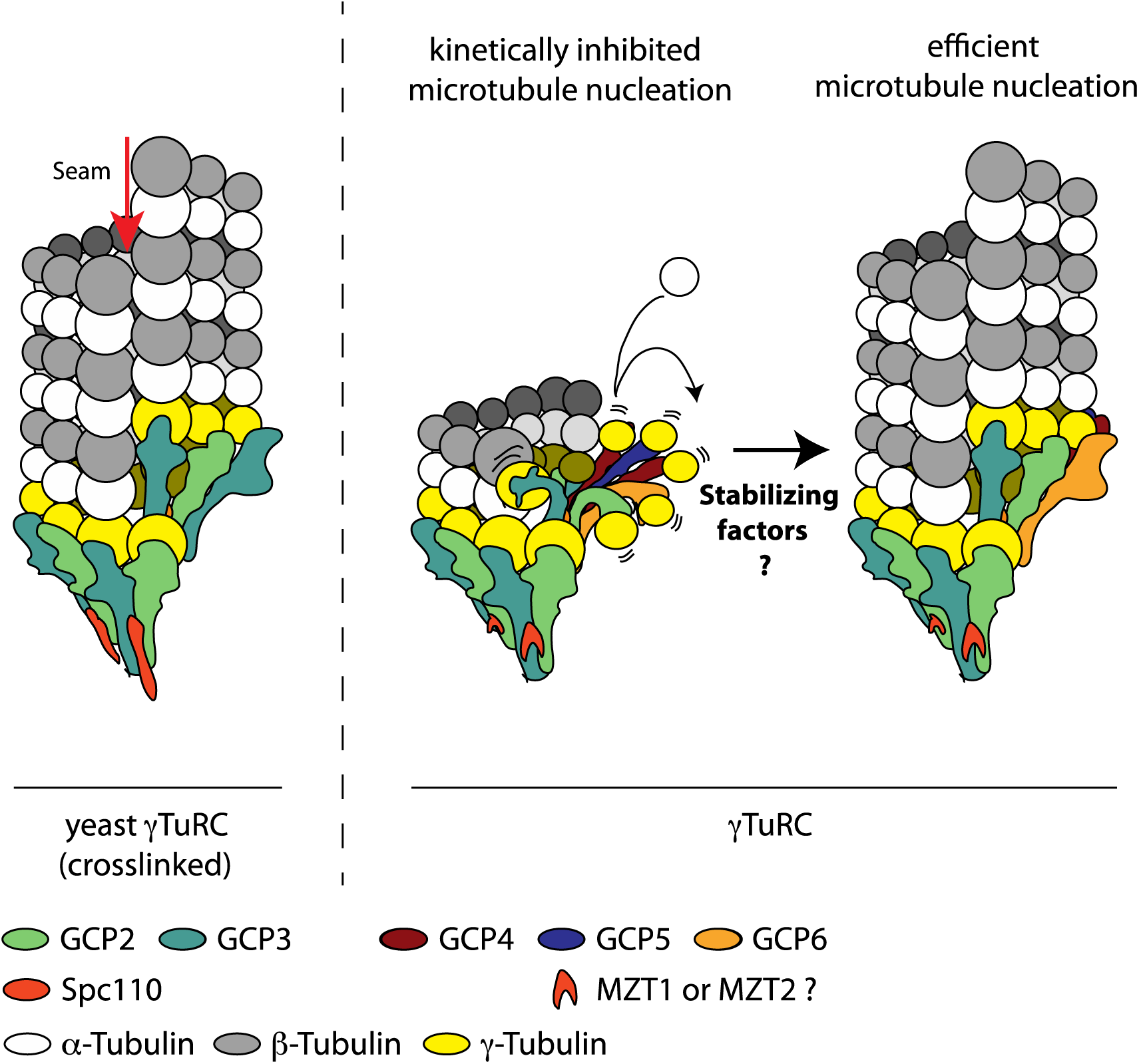
Possible mechanism for efficient microtubule nucleation by γTuRC. Cross-linking of yeast γTuRC has been shown to activate efficient microtubule nucleation by stabilising a closed configuration that recapitulates the 13 protofilament geometry of a microtubule. Our γTuRC structure shows an asymmetric configuration with 4 GCP2, 3 heterodimers positioning γ-tubulin in a closed state competent for nucleation. GCP4, 5, 6 are instead found in an open configuration that is likely to be inactive. Recruitment of GCP stabilising factors likely promotes efficient microtubule nucleation.

## SUPPLEMENTARY TABLES

**Table S1:**
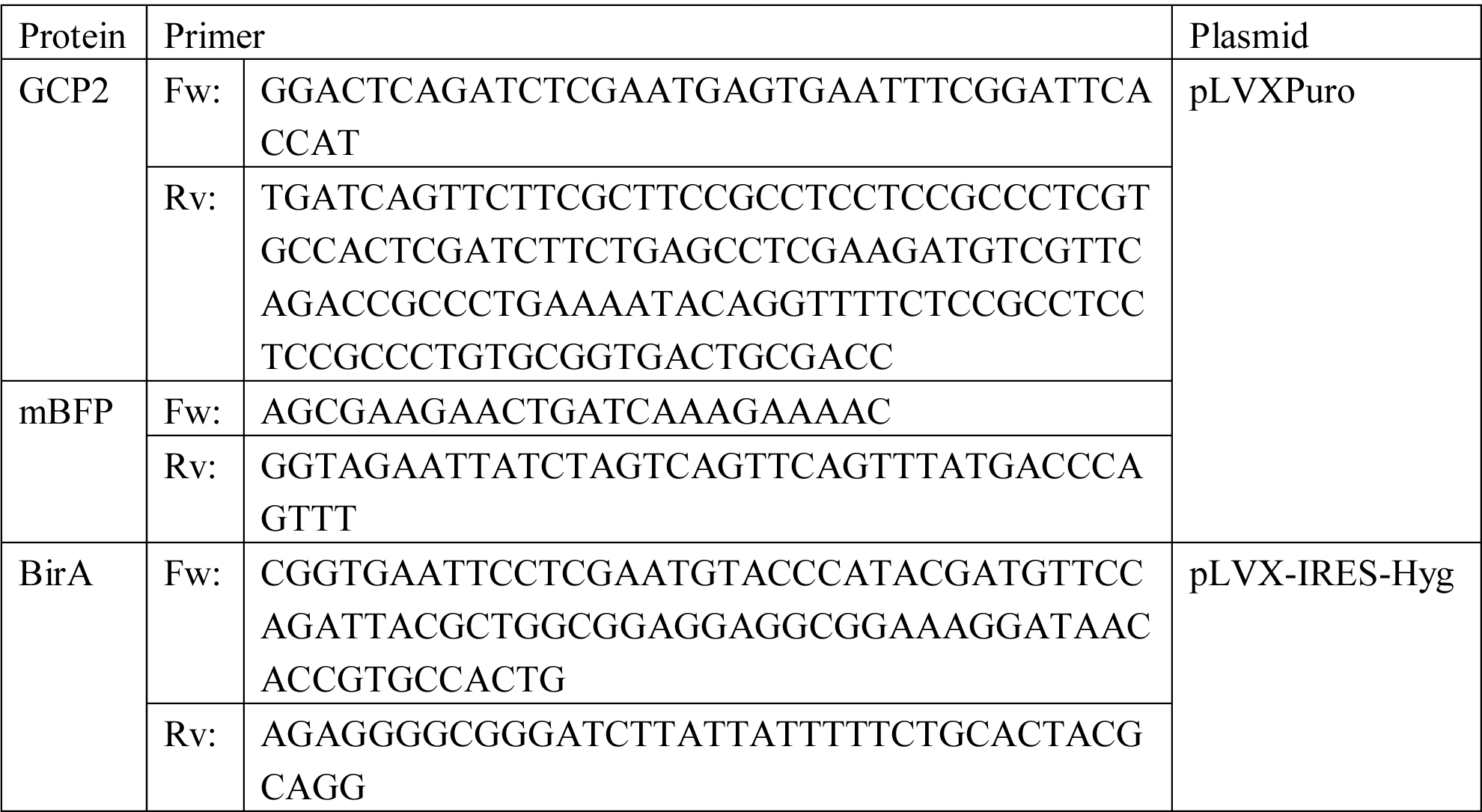
Primers and plamids used for cloning of lentiviral expression constructs.

**Table S2:**
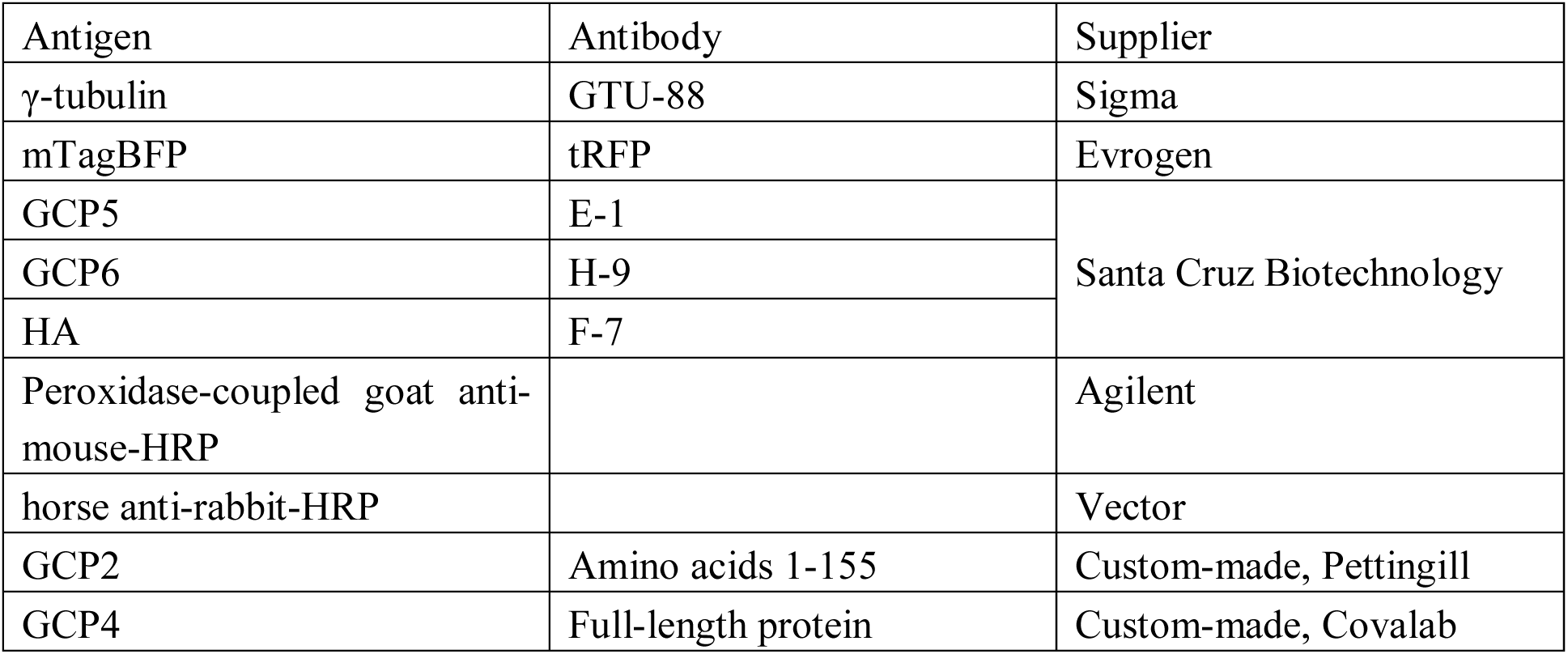
Commercial and custom-made antibodies used for characterization of purified γTuRC by western blotting.

| Antigen | Antibody | Supplier |
| --- | --- | --- |
| γ-tubulin | GTU-88 | Sigma |
| mTagBFP | tRFP | Evrogen |
| GCP5 | E-1 | Santa Cruz Biotechnology |
| GCP6 | H-9 |  |
| HA | F-7 |  |
| Peroxidase-coupled goat anti-mouse-HRP |  | Agilent |
| horse anti-rabbit-HRP |  | Vector |
| GCP2 | Amino acids 1-155 | Custom-made, Pettingill |
| GCP4 | Full-length protein | Custom-made, Covalab |

## MOVIE LEGENDS

**Movie 1. γTuRC nucleated microtubules are capped at their minus ends.** Two microtubules nucleated from a γTuRC surface (373 pM γTuRC used for immobilization) in the presence of 15 μM CF640R-tubulin. The static end is marked with a purple arrow. Time is in seconds. Scale bar is 5 μm. This movie relates to Fig. 2a.

**Movie 2. mGFP-EB3 tracks the growing microtubule plus end of γTuRC nucleated microtubules.** Two microtubules nucleated from a γTuRC surface (373 pM γTuRC used for immobilization) in the presence of 12.5 μM CF640R-tubulin (magenta) and 200 nM mGFP-EB3 (green). mGFP-EB3 tracks the growing microtubule plus-end while the microtubule minus-end is static. Time is in seconds. Scale bar is 5 μm. This movie relates to Fig. 2d.

**Movie 3. Microtubules originate from single immobilized γTuRC molecules.** CF640R-labelled microtubules (magenta) nucleating from single mBFP-labelled γTuRC molecules (cyan, 27 pM γTuRC used for immobilization). Tubulin concentration was 20 μM. Time is in seconds. Scale bar is 2 μm. This movie relates to Fig. 2f.

**Movie 4. Microtubule nucleation efficiency depends on γTuRC surface density.** Microtubule nucleation and growth of CF640R-labelled microtubules from a surface containing an increasing density of immobilized γTuRC (left to right, 47 pM, 187 pM, 373 pM γTuRC used for immobilization). Tubulin concentration was 15 μM. Time is in minutes. Scale bar is 20 μm. This movie relates to Fig. 3a.

**Movie 5. γTuRC microtubule nucleation efficiency depends on tubulin concentration.** Microtubule nucleation and growth of CF640R-labelled microtubules from a surface containing immobilized γTuRC (373 pM used for immobilization) in the presence of increasing concentration of tubulin (from left to right, 7.5 μM, 12.5 μM, 20 μM). Time is in minutes. Scale bar is 20 μm. This movie relates to Fig. 3c.

**Movie 6. chTOG-mGFP synergistically increases γTuRC nucleation efficiency.** Microtubule nucleation and growth of CF640R-labelled microtubules in the presence of γTuRC (left), γTuRC and chTOG-mGFP (middle) or mGFP-chTOG (right). Tubulin concentration was 10 μM (magenta), chTOG-mGFP concentration was 100 nM (green) and 373 pM of γTuRC was used for immobilization. Time is in minutes. Scale bar is 20 μm. This movie relates to Fig. 4a.

**Movie 7. mGFP-TPX2 increases γTuRC microtubule nucleation efficiency in a dose-dependent manner.** Microtubule nucleation and growth of CF640R-labelled microtubules in the presence of γTuRC (left), γTuRC and mGFP-TPX2 (middle) or mGFP-TPX2 (right). Tubulin concentration was 10 μM (magenta), mGFP-TPX2 concentration was 390 nM (green) and 373 pM of γTuRC was used for immobilization. Time is in minutes. Scale bar is 20 μm. This movie relates to Fig. 4e.

**Movie 8. γTuRC structure explains why microtubule nucleation is kinetically inhibited.** The tetramer of GCP2-GCP3 heterodimers (green) follows the geometry of a 13 protofilament microtubule. The GCPs in the asymmetric part (purple) however depart from the microtubule geometry.

